# An automated archival single-nucleus total RNA sequencing platform mapping integrative and retrospective cell atlas of gliomas

**DOI:** 10.1101/2023.11.16.567325

**Authors:** Ziye Xu, Lingchao Chen, Xin Lin, Yuexiao Lyu, Mofei Zhou, Haide Chen, Heng Zhang, Tianyu Zhang, Yu Chen, Yuanzhen Suo, Qian Liang, Zhiyong Qin, Yongcheng Wang

## Abstract

Single-cell RNA sequencing (scRNA-seq) has dramatically transformed biomedical research within laboratory settings. It has been extensively employed to investigate the heterogeneity and plasticity of glioma, the most prevalent brain tumor. However, the clinical diagnosis and treatment of glioma remain complex and challenging, highlighting the need for comprehensive cancer research. Currently available scRNA-seq platforms are insufficient to fulfill the demands posed by large-scale clinical applications. Here, we present an automated high-throughput single-nucleus total RNA sequencing platform, known as AAsnRandom-seq. This platform integrates automated single-nucleus isolation and droplet barcoding systems with the random primer-based scRNA-seq chemistry, designed to accommodate a diverse range of sample types. The performance and versatility of AAsnRandom-seq are validated using over one hundred clinical FFPE and frozen samples. AAsnRandom-seq was applied to archival FFPE samples of various glioma subtypes, including rare clinical samples, and matched primary-recurrent glioblastomas (GBMs), delving into the comprehensive molecular characteristic of glioma at single-cell level. Abundant non-coding RNAs (ncRNAs) with distinct expression profiles within different glioma clusters are detected. Promising recurrence-related targets and pathways are identified from the matched primary-recurrent GBMs. AAsnRandom-seq holds significant application value on large-scale integrative and retrospective clinical research using archived specimens.

## Main

The field of single-cell RNA sequencing (scRNA-seq) technologies has now reached the point where it dramatically transforms medicine research within laboratory settings^1^. Glioma, the most prevalent brain tumor, encompasses a wide range of subtypes and exhibits high heterogeneity at the cellular and molecular levels^2, 3^. Recent studies have utilized single-cell-based technologies^4–6^, especially scRNA-seq^7, 8^, to further corroborated the plasticity of gliomas upon both internal and external factors. Nonetheless, the clinical diagnosis and treatment of glioma remain complex and challenging, highlighting the need for large-scale comprehensive clinical research as a potential avenue for more precise classification, early diagnosis, effective treatment and better outcomes.

While scRNA-seq harbors immense potential for various clinical applications, it faces several challenges and limitations that impede its widespread adoption in clinical contexts. Clinical samples encompass a wide range of materials and sample types, including fresh, fixed and formalin-fixed and paraffin-embedded (FFPE) samples. Excess tumor tissues from surgical and autopsy cases are mainly stored in tissue banks in the form of FFPE samples. However, the capabilities of current scRNA-seq platforms are insufficient for these archival clinical samples. Additionally, in many cases, clinical samples are limited in quantity and quality, especially when obtained through minimally invasive procedures or from specimen banks. Consequently, the sensitivity and coverage of scRNA-seq data put forward higher requirements to ensure the precision and reliability of clinical findings. Moreover, the seamless communication of clinical sample data and patient records among different teams and institutions is fundamental for advancing clinical research. The integration and analysis of scRNA-seq datasets obtained from current various platforms can be time-consuming and introduce potential biases.

Advances in molecular biology, microfluidics, and sequencing technology have given rise to a multitude of sample and library preparation methodologies customized for various scRNA-seq applications. The initial and foremost step in conducting scRNA-seq is the isolation of single cells/nuclei with minimized artifacts and RNA degradation^9^. The gentleMACS™ Dissociator^10^ has emerged as a semi-automated instrument for preparing single-cell suspensions from fresh and fixed material, but it remains inapplicable to FFPE samples. To expand the scope of scRNA-seq application, emerging approaches based on 10X Genomics platform, like 10X Genomics fixed RNA assay, snPATHO-seq^11^, snFFPE-seq^12^ offer innovative avenues for archival samples. However, these oligo(dT)-based or probe-based scRNA-seq methods exhibit limitations in terms of detection sensitivity and transcriptome coverage, as well as a noticeable bias towards 3’-end or targeted genes. In our previous work, we introduced a droplet-based single-nucleus RNA sequencing technology (snRandom-seq) for FFPE tissues, capturing total RNAs with random primers^13^. Several other scRNA-seq methods, such as VASA-seq^14^, MATQ-Drop^15^, RamDA-seq^16^, and scFAST-seq^17^, have similarly adopted random primers or semi-random primers to fully profile the transcriptome of individual cells. However, the cumbersome manual operation and incomplete single-cell/nucleus isolation strategy of current random primer-based scRNA-seq methods make them difficult to apply to large-scale clinical studies.

To overcome these challenges, we have developed an automated high-throughput single-nucleus total RNA sequencing platform, known as AAsnRandom-seq. This platform integrates automated single-nucleus isolation and droplet barcoding systems with the random primer-based scRNA-seq chemistry, designed to accommodate a diverse range of sample types. AAsnRandom-seq enables the automated preparation of single-nucleus suspensions from fresh, frozen, fixed, and FFPE samples. AAsnRandom-seq offers the automated encapsulation of single nucleus, barcoded bead and reagent on a high-throughput scale. The droplet barcoding system is enhanced with artificial intelligence (AI) algorithms that monitor the entire microfluidic process to ensure product quality. We optimize the single-nucleus isolation conditions for each tissue and adopt random primer-based scRNA chemistry to capture total RNAs, including partially degraded RNAs. The performance of AAsnRandom-seq is validated using over one hundred clinical FFPE and frozen samples. Specifically, we apply AAsnRandom-seq on archival FFPE samples of various glioma subtypes, including rare clinical cases, and matched primary-recurrent glioblastomas (GBMs), delving into the comprehensive molecular characteristic of glioma at single-cell level. AAsnRandom-seq detected abundant non-coding RNAs (ncRNAs) with distinct expression profiles within different glioma clusters. Promising recurrence-related targets and pathways are identified from the matched primary-recurrent GBMs. AAsnRandom-seq holds significant application value on archived specimens, expanding the scope of scRNA-seq studies and enabling large-scale integrative and retrospective cell atlas analyses in both clinical and laboratory settings.

## Result

### Overview of the AAsnRamdom-seq platform

We designed an automated platform for single-nucleus total RNA sequencing, comprising four main modules: (1) automated single-nucleus isolation, (2) *in situ* RNA capture, (3) automated droplet barcoding, and (4) sequencing (**Fig. 1a**). The flexible and automated dissociation system of AAsnRamdom-seq is suitable for various sample types, including FFPE, fixed, frozen, and fresh samples. Small slices or grains were taken from the regions of interest in the samples and placed into the automated single-nucleus isolation instrument where they were processed with optimal parameters. Following the completion of this procedure, we obtained a single-nucleus suspension, suitable for subsequent experiments or storage at −80℃. Specifically optimized lysis and digestion conditions were developed for different tissues and sample types (**Supplementary Table 1**). All RNA molecules within each individual single nucleus were captured through the multiple annealing of random primers. To address the challenges posed by very small samples, minimizing batch effects, or reducing reaction costs, an optional pre-indexing strategy was offered during the reverse transcription step. This strategy involved dividing nuclei into different tubes for reverse transcription, each with specific pre-indexed random primers, and subsequently pooling them for further processing. The resulting cDNAs in the reverse transcription step were added with poly(A) tailings, enabling the synthesis of the second DNA strand using barcoded oligo-dT primers. To alleviate operational burden and improve read quality, we developed an automated microfluidic system, aided by AI algorithm, for droplet barcoding. In this system, single nucleus, reaction regent, and barcoded bead were automatically encapsulated into droplet. The generated droplets were identified by the AI model, and the collection process only executes once the microfluidic system was confirmed to be in a stable state.

**Figure 1.**
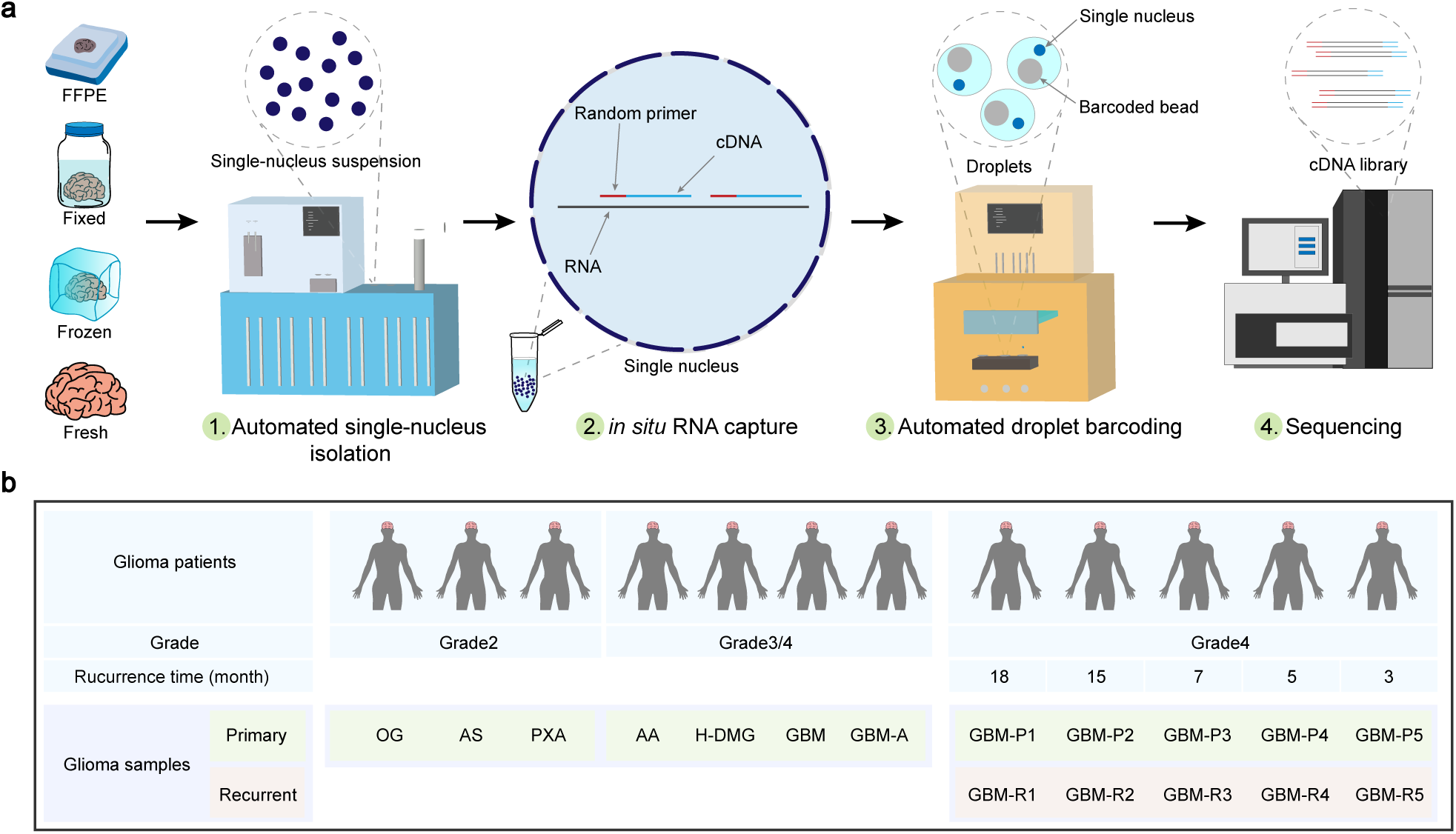
Overview of the AAsnRamdom-seq platform. **a**, The flowchart delineating the AAsnRandom-seq workflow. **b**. The table presents details of glioma FFPE samples performed with AAsnRandom-seq. OG: oligoastrocytoma, AS: astroglioma, PXA: pleomorphic xanthoastrocytoma, AA: anaplastic astrocytoma, H-DMG: high-grade diffuse midline glioma, GBM: glioblastoma, GBM-A: adjacent tissue of glioblastoma, GBM-P: primary glioblastoma, GBM-R: recurrent glioblastoma.

FFPE tissue blocks are the traditional format for biospecimen preservation, and they remain to serve as the most cost-effective means for the long-term storage of tissue specimens. This study focused on the feasibility and robustness of AAsnRandom-seq on clinical FFPE samples. We assessed the performance of AAsnRandom-seq using over one hundred clinical FFPE tissue blocks, which included a diverse array of cancer types such as brain cancer, lung cancer, ovarian cancer and others (**Supplementary Table 1**). We conducted an integrative and retrospective scRNA-seq study on glioma, employing FFPE surgical tumor specimens from various glioma subtypes, including rare clinical cases, and matched primary-recurrent GBMs (**Fig. 1b**). Our primary objectives include the construction of a comprehensive single-nucleus atlas spanning different glioma subtypes, the elucidation of specific expression patterns of non-coding RNAs in gliomas, and the discovery of potential targets associated with tumor processes.

### Automated single-nucleus isolation and droplet barcoding systems

The first challenge encountered in large-scale single-nucleus RNA-seq was the efficient isolation of clear and intact single nucleus, while preserving RNA integrity to the maximum extent possible. To overcome this challenge, we devised an automated single-nucleus isolation system, comprised a user-friendly man-machine interface, a programmable mechanical controller, a temperature controller, and reservoirs for samples and reagents (**Fig. 2a**). Various accessories, including reagent syringes, pipette tips, grinding rods, reaction tubes, collecting tubes, and cell filters, were integrated into the setup (**Supplementary Fig. 1a**). The workflow of this system includes four key stages: paraffin dissolution and rehydration (applicable solely to FFPE samples), lysis, digestion, and filtration (**Supplementary Fig. 1b**). The internal structure and flow pathways of this automated single-nucleus isolation are depicted in **Supplementary Fig. 1c**. Clean and intact single nuclei with a size distribution ranging from 5 to 15 μm were isolated (**Fig. 2b**, **Supplementary Fig. 1d**). The lysis and digestive buffers, as well as the digestion time, were optimized to suit specific tissue types (**Supplementary Table 1**). Solid tissues from parenchymal organs (e.g., liver, muscle, heart) required the strongest lysis and digestive condition. Gastrointestinal tissues (e.g., gastric carcinoma) and immune tissues (e.g., lymphoma) demanded milder conditions.

**Figure 2.**
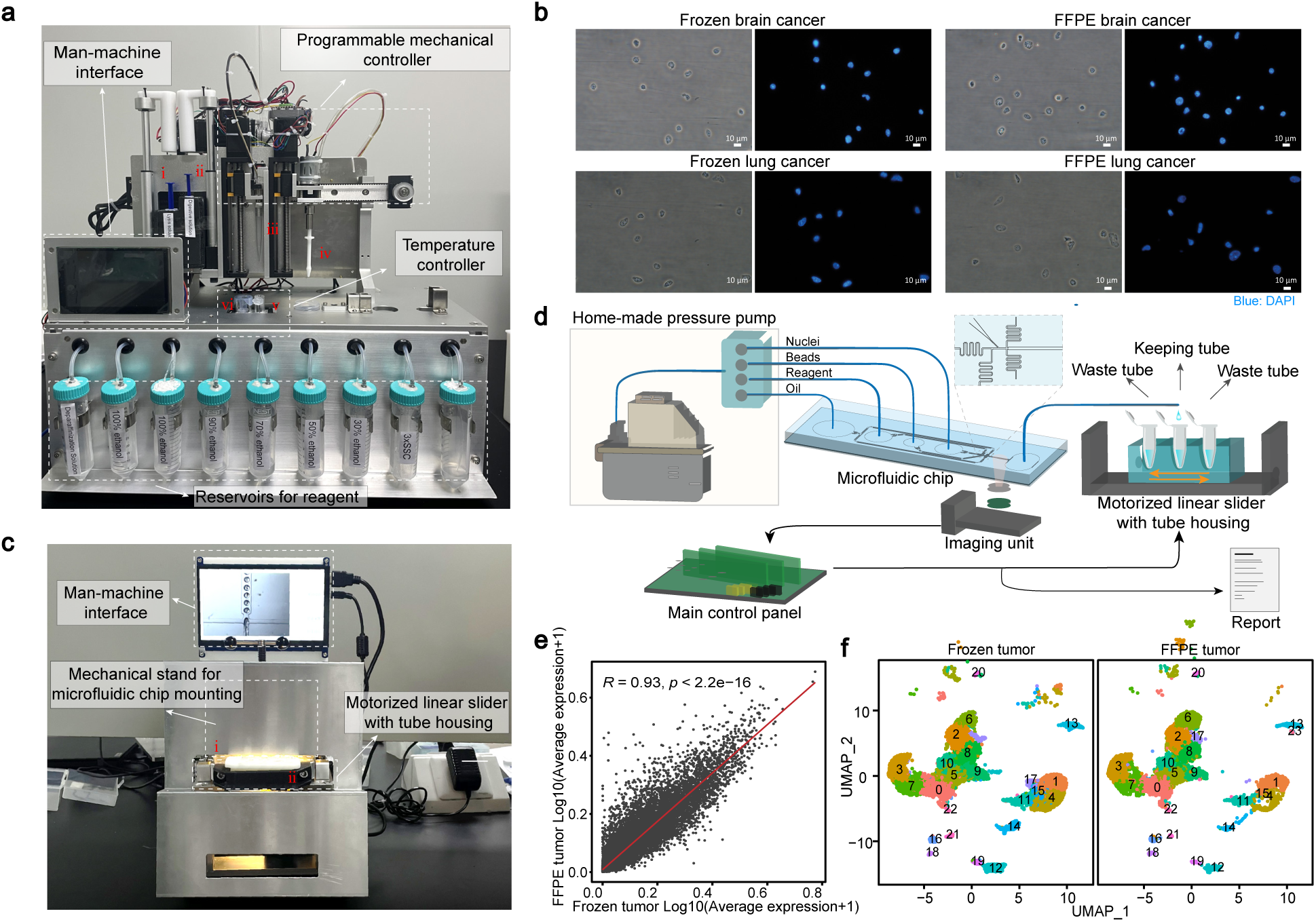
Automated single-nucleus isolation and droplet barcoding systems. **a**, The photograph depicting the arrangement of components within the automated single-nucleus isolation system of AAsnRandom-seq. The corresponding numerical codes for these system accessories are as follows: ⅰ, regent syringe, ⅱ, pipette tip, ⅲ, grinding rod, ⅳ, reaction tube, ⅴ, collecting tube, ⅵ, cell filter. **b**, Microscopical images of nuclei isolated from frozen and FFPE tumor samples from the same brain and lung cancer tissues by AAsnRandom-seq, stained by DAPI. Scale bar, 10 μm. **c**, The photograph depicting the arrangement of various components within the automated droplet barcoding system of AAsnRandom-seq. **d**, A schematic diagram of the automated and AI-assistant droplets collecting system. **e**, The dot plot showing the Pearson’s correlation coefficient (*R*) of the average normalized gene expression levels between the frozen and FFPE tumor samples from the same glioblastomas (GBM) tissue. Each dot corresponds to the average normalized expression level of a gene. The red line indicates the linear regression line. *p* value (*p*) was calculated using a two-sided permutation test. **f**, The integrated UMAP maps generated from the AAsnRandom-seq data of frozen and FFPE tumor samples from the same GBM tissue. Source data are provided as a Source Data file.

We also developed an automated microfluidic system, aided by AI algorithm, to achieve precise encapsulation of a single nucleus and a barcoded bead within each droplet, eliminating the need for intricate manual procedures and expert proficiency (**Fig. 2c**). The configuration of this automated droplet barcoding system includes various components: a home-made pressure pump, a motorized linear slider with tube housing, an imaging unit, a main control panel, and a mechanical stand for microfluidic chip mounting (**Fig. 2d**). The accessories include microfluidic chips and colleting tubes (**Supplementary Fig. 2a, b**). Within this system, mixed reagents, single-nucleus suspension, beads, and oil were directly loaded into the microfluidic chip and subsequently propelled into the channel using a home-made pressure pump to divert the pressure (**Fig. 2d**). Continuous images of droplet generation were captured by a camera, and then processed by AI image recognition technology integrated into the main control panel (**Supplementary Fig. 2c**). By providing microscope images of various droplets for machine learning model training, the system can identify droplets that lack beads, as well as those containing one or more beads (**Supplementary Fig. 2d**). The initial and final stages of droplet generation were found to be particularly prone to producing unstable droplets. To address this, the main control panel selectively adjusts the motorized linear slider with tube housing based on the number of qualified droplets identified, thereby improving the quality of droplet barcoding in AAsnRandom-seq (**Supplementary Fig. 2e**). Additionally, the microfluidic runner can sometimes become inadvertently clogged with unexcepted impurities during the encapsulation process. For quality control, this automated system droplet barcoding system generates a report that assesses the quality of droplets, including a series of parameters, such as droplet counts, diameters, counts of droplets containing 0, 1, or more than 1 bead, and the rate of qualified droplets (**Fig. 2d**).

We validated the feasibility and robustness of AAsnRandom-seq using over one hundred clinical cancer FFPE samples (**Supplementary Table 1**). We next applied it to both FFPE and frozen samples obtained from the same glioma case and revealed a notably high correlation (*R*= 0.93, *p* < 2.2e-16) in the average gene expression levels between FFPE and frozen glioma samples (**Fig. 2e**). Additionally, UMAP plots displaying a substantial overlap in cell clusters between FFPE and frozen glioma samples, offered additional evidence to support the utility of AAsnRandom-seq (**Fig. 2f**).

### Integrative single-nucleus atlas of glioma subtypes

Previous scRNA-seq studies of gliomas have predominantly concentrated on GBM, leaving a gap in our understanding of the comprehensive heterogeneity among various glioma subtypes. In this study, we performed AARamdon-seq on FFPE tumor samples of six glioma subtypes, including the rare clinical cases, as well as one FFPE adjacent tissue sample (**Supplementary Table 2**). The six glioma subtypes includes oligodendroglioma (OG), astrocytoma (AS), pleomorphic xanthoastrocytoma (PXA), glioblastoma (GBM), high-grade diffuse midline gliomas (H-DMG), and anaplastic astrocytoma (AA). Using unsupervised clustering on the integrated AAsnRandom-seq dataset of these FFPE samples, nuclei from these distinct samples were considerably overlapped within the UMAP plot (**Fig. 3a**), and then separated into 37 clusters based on gene expression patterns (**Supplementary Fig. 3a**). According to the expression levles of known markers (**Supplementary Fig. 3b**), each cluster was classified as either glial cells, microglial cells/macrophages, T cells, fibroblasts, excitatory neurons, inhibitory neurons, endothelial cells, pericytes, and proliferating glial cells (**Fig. 3a**). Each tumor subtype contained all these identified cell types (**Supplementary Fig. 3c**). However, the proportions of cellular composition exhibited notable difference between tumor (GBM) and adjacent tissue (GBM-A). The proportions of cellular composition also displayed variations among different glioma subtypes. As expected, glial cells were the most abundant cell type in these gliomas, except in the case of PXA, where fibroblasts constituted 50% of all profiled nuclei. We calculated the top five signature genes of each cell types (**Supplementary Fig. 3d**). Besides the established cell type markers, such as *MBP* for oligodendrocytes and *COL1A1* for fibroblasts, we discovered potential markers for these cell types in gliomas, such as *SLC4A4* for glial cells. Furthermore, we confirmed the elevated expression of PTPRZ1 in glial cells, which was previously considered an oncogene implicated in tumor promotion and invasion in gliomas^18^.

**Figure 3.**
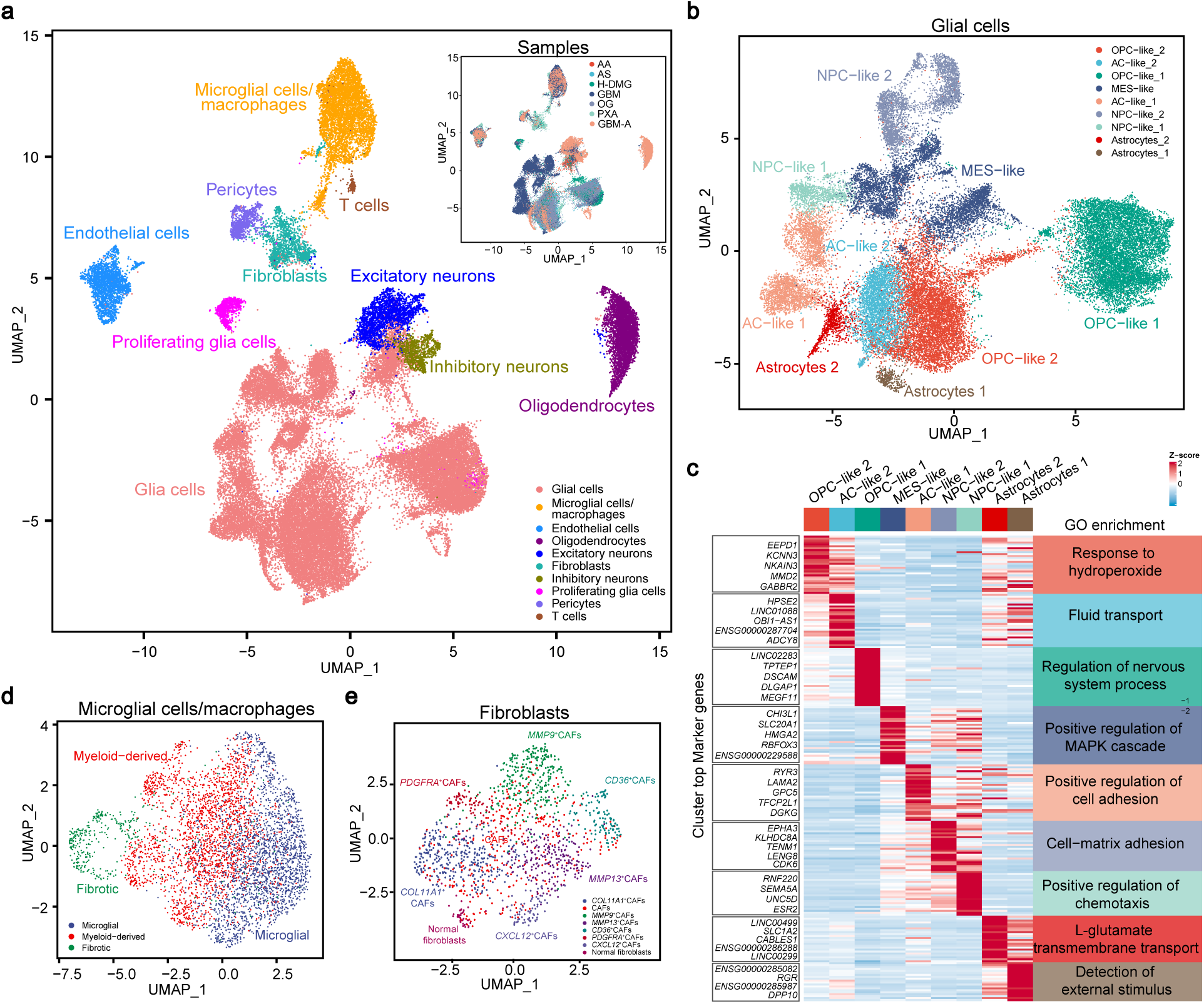
Integrative single-nucleus atlas of glioma subtypes. **a**, UMAP analysis of nuclei isolated from the seven FFPE samples. The big UMAP plot is colored by identified cell types. Ten cell types were identified. The smaller UMAP plot in the upper right corner of Fig. 3a is colored by samples. OG: oligoastrocytoma, PXA: pleomorphic xanthoastrocytoma, AA: anaplastic astrocytoma, AS: astroglioma, H-DMG: high-grade diffuse midline glioma, GBM: glioblastoma, GBM-A: adjacent tissue of glioblastoma**. b**, UMAP analysis of the identified glial cells and colored by identified subclusters. Nine subclusters of glial cells were identified. OPC: oligodendrocyte-progenitor-like, AC-like: astrocyte-like, MES-like: mesenchymal-like, NPC-like: neural-progenitor-like. **c**, A heatmap showing the unique top 5 differentially expressed genes in the nine subclusters of glial cells, ranked by average log2(foldchange). Average gene expression values were scaled and transformed to a scale from −2 to 2. The enriched GO terms of the top 30 differentially expressed genes were shown on the right side of the heatmap. **d**, **e**, UMAP analysis of the identified microglial cells/macrophages (**d**) and fibroblasts (**e**), colored by identified subclusters. CAFs: cancer-associated fibroblasts.

We proceeded by isolating glial cells and performing de novo clustering (**Supplementary Fig. 4a, b).** The nonmalignant glial cells were categorized as astrocytes-1 and astrocytes-2 (**Fig. 3b**). The malignant glial cells were categorized as OPC-like, NPC-like, AC-like, or MES-like cell states, according to the expression levels of established marker genes (**Supplementary Fig. 4c**). Consistent with previous studies^19^, each glioma subtype exhibited a heterogeneous composition of glial cells across these four cell states (**Supplementary Fig. 4d**). Compared to other glioma subtypes, GBM exhibited a higher proportion of NPC-like glial cells (NPC-like 1 and NPC-like 2). These particular subpopulations of glial cells expressed distinct gene expression patterns, including both mRNAs and long non-coding RNAs (lncRNAs) (**Fig. 3c**). *CHI3L1*, a gene previously implicated in supporting tumor growth by influencing the state of glioma stem cells^20^, exhibited high expression in the MES-like subcluster. The lncRNA *OBI1-AS1*, previously recognized as an astrocyte marker with a possible role in glioma recurrence and progression^21^, showed high expression in the AC-like 2 subcluster. We performed Gene Ontology (GO) term enrichment analysis on the top 30 signature genes of these glial cell subpopulations, revealing enrichment in several glioma-related terms^22, 23^ (**Fig. 3c**). Notably, the signature genes of astrocytes-2 subcluster were enriched in L-glutamate transmembrane transport, associated with malignant glioma biology^22^. The signature genes of the MES-like subcluster were enriched in positive regulation of MAPK cascade, associated with glioma invasion and metastasis^23^.

In previous studies, distinct subpopulations of macrophages and fibroblasts localized around tumor cells were identified as tumor-associated macrophages (TAMs)^24^ and cancer-associated fibroblasts (CAFs)^25^, holding promise as potential therapeutic targets. We extracted and conducted de novo clustering on the microglial cells/macrophages and fibroblasts obtained from these gliomas, respectively (**Supplementary Fig. 5a, b, 6a, b**). We classified the microglial cells/macrophages into three groups: microglial, myeloid-derived, and fibrotic macrophages (**Fig. 3d**), based on their expression patterns of specific cell markers (microglial: *CX3CR1*, *P2RY12*, *P2RY13*, and *SELPLG*, myeloid-derived: *CD163*, *TGFBI*, and *F13A1*, fibrotic: *COL1A1*, *COL1A2*, *COL6A3*, and *COL6A2*) (**Supplementary Fig. 5c**). The high-grade gliomas (GDM, H-DMG, and AA) exhibited higher percentages of myeloid-derived macrophages (**Supplementary Fig. 5d**). Notably, PXA displayed a significant presence of fibrotic macrophages, consistent with the previous PXA case reports describing extensive fibrosis^26^. Furthermore, fibroblasts of these gliomas were further subclustered into distinct categories: Normal fibroblasts, CAFs, *COL11A1*^+^CAFs, *MMP9*^+^CAFs, *MMP13*^+^CAFs, *CD36*^+^CAFs, *PDGFRA*^+^CAFs, and *CXCL12*^+^CAFs (**Fig. 3e**), based on the expression patterns of signature genes (**Supplementary Fig. 6c**). Notably, a high percentage of *COL11A1*^+^CAFs was present in PXA (**Supplementary Fig. 6d**), which was consistent with the macrophage constitution of PXA. Subpopulations associated with invasion and metastasis, including *MMP9*^+^CAFs, *MMP13*^+^CAFs, and *CXCL12*^+^CAFs, were mainly present in PXA and H-DMG. Additionally, CellChat analysis of the cell subclusters within PXA sample highlighted a strong interaction of between *PDGFRA*^+^CAFs and fibrotic macrophages (**Supplementary Fig. 6e**).

### Numerous ncRNAs exhibited specific expression within distinct glioma clusters

Most mainstream scRNA-seq studies, relied on the prevalent poly(A)-based RNA capture strategy, only targeted polyadenylated mRNAs. However, this strategy inherently presents limitations in capturing a comprehensive transcriptome, notably missing numerous crucial regulatory RNAs that lack poly(A)-tails. Multiple studies have indicated that non-coding RNAs (ncRNAs) play critical roles in various biological processes of glioma initiation and progression^27^. In our analysis, AAsnRandom-seq demonstrated its proficiency in detecting a lot of ncRNAs, including regulatory ncRNAs (lncRNAs and miRNAs) and constitutive ncRNAs (snRNAs and snoRNAs) across all FFPE glioma samples (**Supplementary 7a**). The lncRNAs detected by AAsnRandom-seq included over 6,500 unannotated and over 3,500 annotated lncRNA (**Fig. 4a**). To further characterize these ncRNAs within gliomas, we extracted the expression matrix of ncRNAs from the merged datasets of these FFPE samples and performed a finding-marker analysis to identify signature ncRNAs across various cell types (**Fig. 4b**). All these signature ncRNAs, including *MIR9-1HG* (in glial cells and proliferating glia), *OBI1-AS1*, and *LINC01088* (in glial cells), *LINC01572* (in proliferating glia), *LINC01608*, *LINC00844*, *LINC00639*, and *LINC01170* (in oligodendrocytes), *LINC01374*, *MIR646HG*, and *DLEU1* (in macrophages), *MIR3667HG* (in macrophages and T cells), *LINC00861* (in T cells), *LINC01250* and *MIR124-1HG* (in excitatory neurons and inhibitory neurons), *JARID2-DT* (in excitatory neurons), *DLX6-AS1* (in inhibitory neurons), *TEX41* and *TTTY14* (in fibroblasts), and *CARMN* (in pericytes), were found to be lncRNA genes. These lncRNAs may hold signature roles in identifying specific glioma cell types.

**Figure 4.**
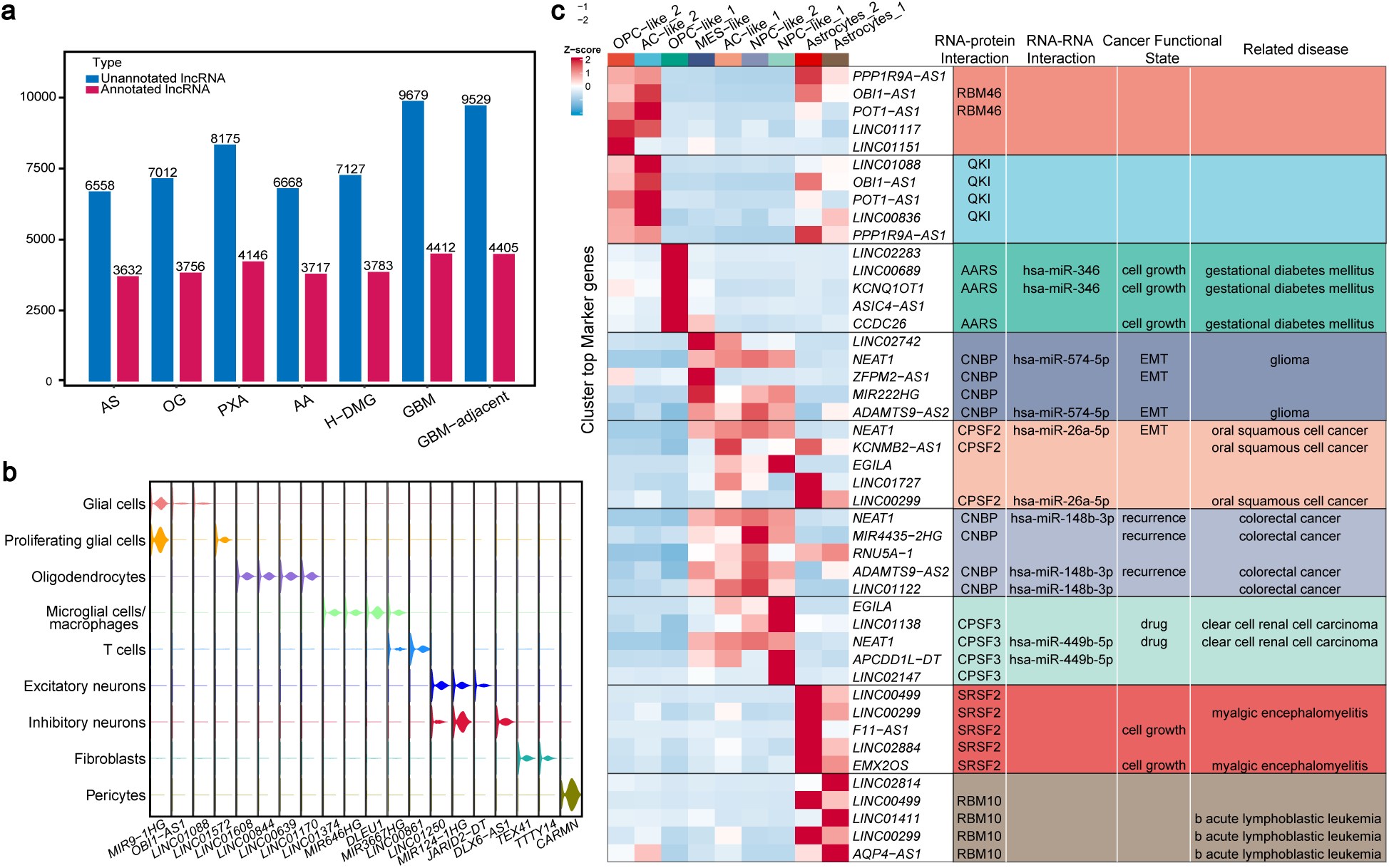
Numerous ncRNAs exhibited specific expression within distinct glioma clusters. **a**, A bar graph showing the numbers of unannotated and annotated lncRNAs detected in the seven FFPE samples by AAsnRandom-seq. **b**, Violin plot of the specifically expressed ncRNAs in different cell types of gliomas. **c**, A heatmap showing the top 5 signature ncRNAs in the nine subclusters of glial cells, ranked by average log2 (foldchange). Average gene expression values were scaled and transformed to a range from −2 to 2. On the right side of the heatmap, information regarding the interacted RNA, protein, cancer function state, and related diseases of these top 5 signature ncRNAs were provided. This information was collected from LncSEA2.0 (http://bio.liclab.net/LncSEA/index.php). Source data are provided as a Source Data file.

We proceeded to focus on the expression matrix of ncRNAs within glial cells and identified signature ncRNAs among the distinct glial cell subpopulations (**Fig. 4c**). Within this set of ncRNAs, we observed several that have been previously implicated as promoters of glioma progression, such as *LINC02283*^28^, *LINC01088*^29^, *LINC00689*^30^ and *NEAT1*^31^. Particularly, *NEAT1* exhibited high expression in the malignant subclusters mainly found in GBM (MES-like, AC-like 1, NPC-like 1, and NPC-like 2). Consistently, patients with GBM exhibiting high NEAT1 expression displayed reduced survival times (Overall Survival: *P* = 0.085, Disease Free Survival: *P*= 0.045) (**Supplementary Fig. 7b, c**). Furthermore, several protein-coding genes associated with these ncRNAs, such as the neighboring coding gene (*SLC7A11*) of signature lncRNA *LINC00499* and the gene (*EMX2*) on opposite strand transcript of signature lncRNA *EMX2OS*, had been previously reported to be associated with glioma^32, 33^. Moreover, we collected the information about RNA-RNA/Protein interactions, cancer function states, and related diseases of these signature ncRNAs (**Fig. 4c**). Notably, the signature ncRNAs of AC-like 2 (*LINC01088*, *OBI1-AS1*, *POT1-AS1* and *LINC00836*) were enriched in the interaction with protein QKI, which plays a regulatory role in glioma stem cell (GSC) stemness^34^. Likewise, the signature ncRNAs of OPC-like 1 (*LINC00689*, and *KCNQ10T1*) and NPC-like 1 clusters (*NEAT1* and *APCDD1L-DT*) were enriched in the interaction with RNAs hsa-miR-346 and hsa-miR-449b-5p, respectively, both of which regulate glioma growth^35, 36^. These signature ncRNAs were enriched in various cancer function states. For example, the signature ncRNAs of OPC-like 1 (*LINC00689*, *KCNQ10T1*, and *CCDC26*) and Astrocytes 2 clusters (*F11-AS1* and *EMX2OS*) were enriched in cell growth. The signature ncRNAs of MES-like (*NEAT1*, *ZFPM2-AS1*, and *ADAMTS9-AS2*) were enriched in epithelial-mesenchymal transition (EMT), consistent with the mesenchymal-related mRNA gene expression signature of MES-like subcluster. These signature ncRNAs were previously reported to be related with various diseases. Notably, the specific-expressed ncRNAs of MES-like (*NEAT1* and *ADAMTS9-AS2*) are related with glioma. Accordingly, these signature ncRNAs detected by AAsnRandom-seq might exert diverse functions at different stages of glial cells and could serve as therapeutic targets for gliomas.

### Retrospective atlas of matched primary-recurrent glioblastomas

Conventional scRNA-seq technologies encounter limitations when attempting to utilize matched primary-recurrent samples from the same patients for longitudinal clinical study^37^. This limitation can potentially result in the omission of crucial information regarding tumor progression. AAsnRandom-seq showed high applicability on archival samples, especially the FFPE samples. We applied AAsnRandom-seq to five pairs of matched primary-recurrent FFPE specimens from GBM patients with different recurrence time (Patient G7: 18 months, Patient G8: 15 months, Patient G9: 7 months, Patient G10: 5 months, Patient G11: 3 months) (**Supplementary Table 2**). We followed the standard scRNA-seq analysis process on the integrated AAsnRandom-seq data to construct a retrospective atlas of GBMs. Nuclei from primary and recurrent FFPE specimens overlapped well on the UMAP plots (**Supplementary Fig. 8a, b**) and clustered into distinct cell clusters (**Supplementary Fig. 8c**). Main cell types of gliomas, including glial cells, microglial cells/macrophages, proliferative glia, oligodendrocytes, endothelial cells, pericytes, fibroblasts, and T cells, were identified in all primary and recurrent samples using known markers (**Fig. 5a, Supplementary Fig. 8d**). Notably, the proportion of oligodendrocytes was significantly higher in recurrent sample R1, which had a longer recurrence time (18 months) (**Supplementary Fig. 8e**). An increase in macrophages and T-cell infiltration is often observed in recurrent GBM^38, 39^. We found that the proportions of immune cells, including microglial cells/macrophages and T cells, were increased in the recurrent group, except for R2, which had a relatively long recurrence time (15 months) (**Fig. 5b**). Differences in the changes of oligodendrocyte and immune cell proportions in these recurrent samples might be associated with the recurrence time and individual patient differences. The proportions of microglial cells/macrophages in primary samples with relatively short recurrence time (7, 5, and 3 months) were higher than that in primary samples with longer recurrence time (18 and 15 months) (**Supplementary Fig. 8e**), suggesting that the proportion of microglial cells/macrophages might be correlated with recurrence time.

**Figure 5.**
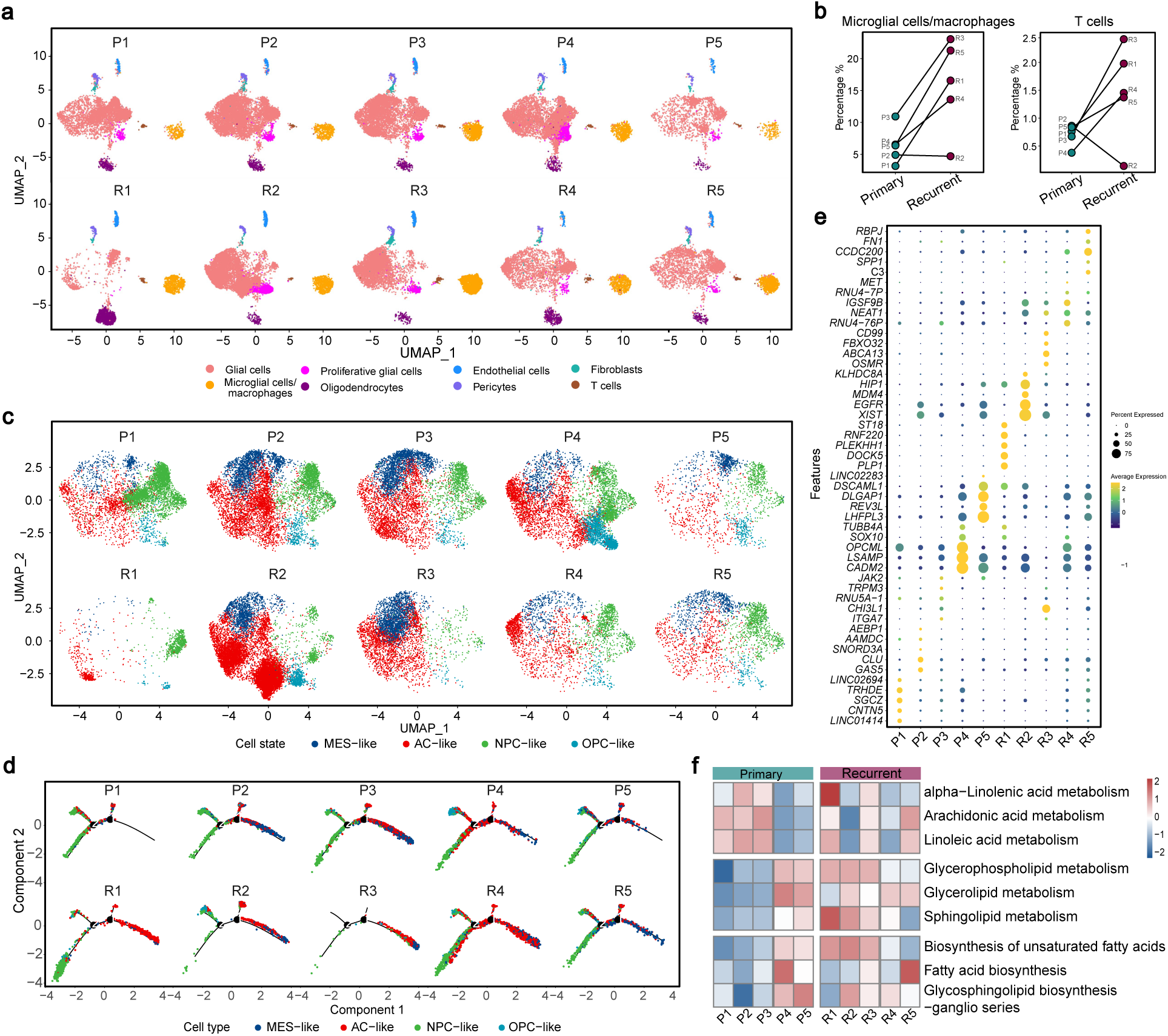
Molecular characteristics of matched primary-recurrent glioblastomas. **a**, Separated UMAP plots of nuclei from five pairs of matched FFPE samples of primary-recurrent GBM cases. UMAP plots are colored by identified cell types. P1-P5: primary GBM sample 1-5. R1-R5: recurrent GBM sample 1-5. **b**, The proportions of microglial cells/macrophages and T cells in primary and recurrent samples. Left panel: microglial cells/macrophages, right panel: T cells. **c**, Separated UMAP plots of glial cells from primary and recurrent GBM samples. UMAP plots are colored by glial cell states. **d**, Separated trajectories of glial cells from primary and recurrent GBM samples generated by monocle analysis and colored by glial cell states. **e**, Dot plot showing the unique top 5 signature genes of OPC-like glial cells among different primary and recurrent GBM samples. **f**. Heatmap showing the expression scores of lipid metabolism pathways in OPC-like glial cells of different primary and recurrent GBM samples. Source data are provided as a Source Data file.

Next, we performed de novo unsupervised clustering of glial cells (**Supplementary Fig. 9a**-c). The four main glial cell states, including MEC-like, AC-like, OPC-like, and NPC-like cluster, were roughly identified according to the previously identified highly expressed genes^19^ (**Fig. 5c, Supplementary Fig. 9d**). The proportions of these four glial subpopulations were quite different among individual patients (**Supplementary Fig. 9e**). Group preference analysis showed that NPC-like and OPC-like cluster were more concentrated in primary samples, while AC-like cluster were more concentrated in recurrent samples (**Supplementary Fig. 9f**). We reconstruct the pseudo-temporal trajectory inference of all glial cells from primary and recurrent GBM using monocle analysis (**Supplementary Fig. 9g**). The glial cells trajectory yielded five developmental hierarchies (State 1-5) where the NPC-like cluster mainly located at the beginning of cell evolution map and the AC-like and MES-like clusters mainly located at the endpoint (**Fig. 5d, Supplementary Fig. 9h**), consistent with the previously reported the transition direction along the NPC/OPC-AC-MES axis^40^. Notably, OPC-like cluster mainly located at a branch of the starting point (state 5), especially in the primary samples P4 and P5, which had relatively shorter recurrence time. Previous studies have shown that OPC-like cells exhibit greater proliferation and tumor-propagating potential^41^. Combined with the higher proportion of OPC-like cluster in P4 and P5 (**Supplementary Fig. 9e**), we speculate that OPC-like glial cells may be associated with GBM recurrence and deserves more attention for therapeutic purposes. We subsequently identified the signature genes of OPC-like cells in these primary and recurrent samples and discovered several glioma-associated genes (*LHFPL3*^42^, *DLGAP1*^43^, and *LINC02283*^28^) highly expressed in the primary samples P4 and P5 (**Fig. 5e**). The oncogenic lncRNA *LINC02283* was also identified as a specific-expressed ncRNA among the glial subpopulations in the above integrative single-nucleus atlas of glioma subtypes (**Fig. 4c**), suggesting that *LINC02283* may be a critical regulator in glioma biology. Lipid metabolism is reported to be significantly dysregulated in gliomas^44^. We evaluated scores of lipid metabolic pathways in OPC-like glial cells from different samples and found that these lipid metabolic pathways were enhanced in recurrent samples (**Fig. 5f**). Interestingly, unsaturated fatty acid (alpha-linolenic acid, arachidonic acid, and linoleic acid) metabolism was depleted in the primary samples (P4 and P5) with relative shorter recurrence time. The glycerophospholipid, glycerolipid, and sphingolipid metabolism in P4 and P5 were stronger than that in primary samples with longer recurrence time. The biosynthesis pathways, including biosynthesis of unsaturated fatty acids, fatty acid biosynthesis, and glycosphingolipid biosynthesis-ganglio series were also stronger in P4 and P5. These results suggested that more accumulation of unsaturated fatty acids in the OPC-like glial cells might be positively associated with GBM recurrence.

## Discussion

We developed an automated platform, AAsnRandom-seq, for high-throughput single-nucleus total RNA sequencing. The automated single-nucleus isolation system within AAsnRandom-seq was tailored to various clinical archival samples, with a particular focus on FFPE samples. By automating the scRNA-seq platform and incorporating AI image recognition technology, we streamlined the complex processes involving molecular biology and microfluidics in scRNA-seq and controlled the encapsulation quality, thereby enhancing efficiency, accuracy, reproducibility, and adaptability. By applying AAsnRandom-seq to diverse glioma subtypes and matched primary-recurrent GBMs, we anticipate the potential for delivering novel insights into clinical studies related to cancer progression, heterogeneity, and therapeutic responses in gliomas.

AAsnRandom-seq, employing random primers, has successfully detected a rich array of signature genes, especially ncRNAs, in different glioma subtypes and primary-recurrent GBMs. Many of these signature genes have previously been recognized as critical contributors to tumorigenesis, metastasis, and therapy resistance, as established by conventional molecular biology approaches. AAsnRandom-seq provides a powerful tool to unraveling the functional roles and dynamics of tumor-related genes at single-cell resolution, offering insights that were previously elusive. Our application of AAsnRandom-seq to over one hundred clinical cancer FFPE tissues has generated scRNA-seq datasets encompassing total RNAs information. We are extending the application of AAsnRandom-seq to more types of human cancers. Integration of these scRNA-seq datasets with clinical and basic research data is also currently underway, which will facilitate the development of a scRNA-seq database containing single-cell total RNAs expression maps across various human cancers. Additionally, the data preprocessing and traditional analysis of scRNA-seq data are time-consuming, energy-draining, and easy to introduce human error. Many researchers have reported automated pipelines for comparative analysis of scRNA-seq datasets^45, 46^, which can be integrated into AAsnRandom-seq platform to prompt the widespread applications of this large-scale and comprehensive scRNA-seq database across a wide range of scientific disciplines.

Clinical patient-derived repositories serve as valuable resources for investigating diseases, elucidating molecular mechanisms, and developing personalized treatment strategies. A recent advancement introduced a high-throughput single-cell DNA-seq method designed specifically for archival FFPE samples^47^, recognizing the substantial value of clinical FFPE samples in medical research and healthcare. Over the years, millions of FFPE specimens have been preserved, often paired with detailed pathological and clinical documentation, making them readily accessible for the study of virtually any disease^48^. It is both important and practicable to assess the generalizability of the biological findings presented in this study and to further explore glioma progression in larger cohorts of FFPE glioma samples. Meanwhile, it is essential to conduct molecular biology experiments on more extensive collections of FFPE glioma samples to further validate the molecular characteristics and potential therapeutic targets of glioma, such as the lncRNA *LINC02283*. The optimized single-nucleus isolation system and random primer-based RNA capture strategy within AAsnRandom-seq render it highly versatile, accommodating various sample types, including fresh, frozen, fixed and FFPE tissues. We are actively exploring opportunities to broaden the applicability of AAsnRandom-seq to encompass samples such as blood, saliva, faeces, and even microbe, thereby extending diverse clinical application scenarios. This rapid cancer data accumulation demands an inevitable trend of experimental instruments automating. More advanced technologies, such as robotics, artificial intelligence, and machine learning, are integrating into experimental processes. We are committed to the development of a fully-automated, sample-in-result-out scRNA-seq platform, aiming to facilitate its utilization in screening, diagnosis, treatment monitoring, and prognosis evaluation across a wide spectrum of conditions, such as metabolic diseases, coagulation system diseases, pregnancy, infectious diseases, emergency medicine and various other clinical contexts.

Recent years have seen an evolving concept of ‘big data’ in clinic catalyzed by breakthroughs in high-throughput technologies. A fully automated and integrated high-throughput single cell/nucleus total RNA sequencing platform, including kit, instrument, and software, combining utilization of databases, holds the potential to emerge as a vital clinical tool for cancer diagnosis and treatment.

## Methods

### Experimental Model

The collection of human samples and research conducted in this study were approved by the Ethics Committee of Huashan Hospital, Fudan University (approval numbers: KY2022-762) and the Research Ethics Committee of the First Affiliated Hospital, Zhejiang University School of Medicine (approval numbers: IIT20220893A). Clinical FFPE and frozen glioma samples were provided by Huashan Hospital, Fudan University. The other archival clinical FFPE and frozen tumor samples were provided by the First Affiliated Hospital, Zhejiang University School of Medicine.

### Automated single-nucleus isolation

We designed the automated single-nucleus isolation instrument and customized it in collaboration with M20 Genomics Company. The setup and accessories of the automated single-nucleus isolation system are shown in **Fig. 2a** and **Supplementary Fig. 1a**. The setup includes a man-machine interface, a programmable mechanical controller, a temperature controller, and reservoirs for samples and reagents. The accompanying accessories include syringes for reagents, pipette tips, grinding rods, reaction and collection tubes, and cell filters. For FFPE samples, dewaxing and rehydration reagents were introduced into the tube and incubated with vibration, followed by sequential removal. To ensure complete paraffin removal, the dewaxing step was repeated once or twice, as dictated by the specific sample requirements. The tissue was quickly and completely grinded in the presence of lysis buffer under a low temperature (∼4℃). Then, tissue was dissociated by digestive enzymes (Collagenase under ∼37℃, Protein K under ∼50℃). The resulting mixture of dissociated tissue and buffer was filtered, and the single-nucleus suspension was collected. The detailed internal structure and the flow directions of the automated single-nucleus isolation system were illustrated in **Fig. 2c**. The dewaxing and rehydration regents were sucked and injected into the injector tube in turn by the syringe device. All the waste liquid was siphoned off by the pipette tip at the end of each reaction and stored in the waste liquid tank. The lysis buffer was injected by the syringe device into the injector tube. After lysis and digestion, the mixture was transferred to the cell filter above the collecting tube by the pipette tip.

The lysis buffer, digestive enzyme, digestive time of various tissue types were provided in **Supplementary Table 1**. The high lysis buffer contained 1X PBS buffer, 0.2% Nonidet(R)P-40 (NP-40, Sangon Biotech, Cat # A600385), and 1 U/μL RNase Inhibitor. The medium lysis buffer contained 1X PBS buffer, 0.1% TritonX-100 (Sangon Biotech, Cat # A600198), and 1 U/μL RNase Inhibitor (Yeasen Biotechnology, Cat # 10610ES03). The low lysis buffer contained 1X PBS buffer, 0.1% Tween-20 (Sangon Biotech, Cat # A600560), and 1 U/μL RNase Inhibitor. Digestive enzymes, including 1 mg/mL Proteinase K (Sangun Biotech, Cat # A610451) or 1 mg/mL Collagenase I (Gibco, Cat # 17100017) were used.

### *In situ* DNA block

The single nucleus suspension was assessed and quantified through DAPI staining under a fluorescent microscope. Block primers, in accordance with the sequence provided in our previous work^13^, were ordered from Sangon Biotech company (China). The reaction mixture was prepared as follows: 100,000∼1000,000 nuclei in 25.5 µL of PBS, 5 µL of 10 µM block primers, 2 µL of DNA Polymerase (M20 Genomics, Cat # R20123124), 10 µL of 5X DNA polymerization buffer, 5 µL of 100 mM dNTP, 2.5 µL of RNase Inhibitor. This mixture was incubated at 37℃ for 30 minutes. Then, nuclei were subjected to three washes with PBST (1X PBS with 0.05% T-ween 20) to eliminate any residual primers and reagents.

### *In situ* reverse transcription

Following the *in situ* DNA block, we proceeded with *in situ* reverse transcription. Random primers were ordered from Sangon Biotech company (China) according the sequence provided in our previous work^13^. The reaction mixture was composed of 100,000∼1000,000 nuclei in 27.5 µL of PBS, 5 µL of 10 µM random primers, 2.5 µL of Reverse Transcriptase (M20 Genomics, Cat # R20123124), 10 µL of 5X reverse transcription buffer, 2.5 µL of 100 mM dNTP, 2.5 µL of RNase Inhibitor. The reaction mix was incubated with twelve cycles of multiple annealing ramping from 8℃ to 42℃ and 30 min at 42℃ on a thermocycler. Then, nuclei were washed with PBST three times to wash away the residual random primers and regents.

### dA tailing

dA tailing was performed after *in situ* reverse transcription. The following reaction mix was prepared: 100,000∼1000,000 nuclei in 39 µL PBS, 5 µL 10X TdT reaction buffer, 0.5 µL TdT enzyme (NEB, Cat # M0315S), 0.5 µL 100mM dATP (NEB, Cat # N0440S), 5 µL CoCl_2_, and incubated at 37 °C for 30 minutes. Then, nuclei were washed with PBST three times to wash away the residual reagents.

### Design and fabrication of the microfluidic device

The microfluidic chip shown in **Supplementary Fig. 2a** was designed using a computer-aided design software AutoCAD (2021, AutoDESK, USA). The established protocols^49^ was employed for fabricating the polydimethylsiloxane (PDMS) microfluidic chip, with a channel depth of 50 μm. Molds for a microfluidic device were made using a photolithographic approach, consisting of centrifugal coating and modeling the SU-8. Silicon molds were employed for casting PDMS (Sylgard-184) to fabricate microfluidic devices.

### Second strand cDNA synthesis

The morphology of nuclei after *in situ* reactions was observed by optical microscope. Nuclei were counted and diluted with a 30% density gradient solution. Nuclei, 2X DNA extension reaction mix and barcoded beads (M20 Genomics, Cat # R20123124) were encapsulated into droplets using the microfluidic platform. Then, the emulsions were incubated at 37 °C for 1 hour, 50 °C 30 min, 60 °C 30 min, and 75 °C 20 min. After the barcoding reaction, droplets were broken by mixing with PFO buffer. The aqueous phase was taken out and purified by Ampure XP beads (Beckmen, Cat #A63881). PCR primers (Primer1 and Primer2) were ordered from Sangon Biotech company (China) according the sequence provided in our previous work^13^. PCR was performed to amplify the purified product using Primer1 and Primer2 primers. The amplified product was purified by Ampure XP beads and quantified by Qubit.

### AI-assistant droplets collecting

As illustrated in **Fig. 2c** and **2d**, we developed an AI-assistant droplet collecting system consisting of a home-made pressure pump, a motorized linear slider with tube housing, an imaging unit, a main control panel, and a mechanical stand for microfluidic chip mounting. The home-made pressure pump was integrated within the AI-assistant droplet collecting system and capable of supplying at least 4 independent pneumatic pressure sources with the range of 0∼40 kPa and stability of <0.02 kPa. The imaging unit consists of a LED lighting source, a CMOS camera (MER2-U3, Daheng Imaging) and a 4X lens and was mounted underneath the droplet generation region of the microfluidic chip. Once the experiment was triggered, droplet generation within the microfluidic chip was real-time monitored and analyzed. The captured images for droplet generation were sent to the main control panel for processing and video clips were shown to the experiment operator via the man-machine interface. At the beginning and ending stages, unqualified droplet series were discarded to the waste tube. Once droplet quality was confirmed by the control panel, the motorized liner slider was switched and therefore qualified droplets were collected to the keeping tube. After the experiment was completed, a summary report including a series of parameters, such as droplet counts, diameters and etc., will be shown in the man-machine interface and/or PDF version was available for download.

### Real-time image processing

Dataset including more than 5,000 droplet generation images was collected from our previous experiment and annotated manually. The computer vision model was then trained using YOLOv5 and verified using 1000+ additional experimental images. During the experimental course, real-time droplet generation was recorded by the imaging unit and then droplet quality was analyzed and processed by control panel using the trained model.

### Library preparation

The sequencing library was constructed according to VAHTS Universal DNA Library Prep Kit (Vazyme, Cat #ND607-01) for Illumina V3. About 50 ng of DNA fragments were used to construct sequencing library. The input-DNA and final library were quantified by Qubit2.0 (Life Technologies). The fragment sizes of input DNA and final library were measured with Qsep100™ DNA Fragment Analyzer (BIOptic). The DNA fragments were purified and selected using AMPure XP beads. Sequencing was performed using the NovaSeq 6000 and S4 Reagent Kit with paired end reads of 150.

### Data Analysis

#### Preprocessing of AAsnRandom-seq data

First, primer sequences and extra bases generated by the dA-tailing step were trimmed in raw sequencing data. Then for each Read1, we extracted UMI (8 nts) and cell-specific barcode (30 nts) and merged sequenced barcodes that can be uniquely assigned to the same accepted barcode with a Hamming distance of 2 nts or less. Read2 was used to generate the gene expression matrix by the STARsolo module in STAR (2.7.10a) with reasonable parameters. To determine the number of nuclei in each sample, we plotted the scattergram of log10(genes) for each possible barcode and used the position of the minimum with the highest value of log10(genes) as the threshold: only barcodes with the number of genes above this threshold were used for downstream analysis.

#### Clustering and Downstream Analysis

The gene expression matrix was generated after barcode filtering and removal of mitochondrial RNAs and ribosomal RNAs. The analysis and visualization of AAsnRandom-seq data were conducted using the Seurat 3 toolkit^50^ within Rstudio (4.2.1), which encompassed a range of processes: preprocessing, integration, visualization, clustering, cell type identification, and differential expression testing. Genes detected in fewer than 3 nuclei were filtered out. The following thresholds were used for nuclei-level filtering: nCount_RNA< CountThresh & nFeature_RNA>200. CountThresh = mean(nCount_RNA)+2*sd(nCount_RNA). For the integration of AasnRandom-seq datasets, counts were normalized and scaled in Seurat. The integration was executed using the Harmony package^51^ in R. Integrations were performed across the FFPE/fresh comparison samples, tumor subtypes, and primary/recurrent comparison samples, respectively. Within each sample, 2000 anchors were identified, and the integration of AasnRandom-seq datasets was realized through the IntegrateData function, utilizing 20 dimensions^52^. To construct integrated datasets, the shared nearest neighbor (SNN) graph was created by conducting principal component analysis (PCA), followed by the application of FindNeighbors using 30 principal components. Clusters were subsequently delineated using the FindClusters function with a resolution of 1. The visualization of clusters used UMAP of the principal components, as implemented in Seurat. The cell type identification for each cluster was accomplished manually using a published set of marker genes. Marker genes were identified using the FindAllMarkers function within Seurat. The resultant marker genes matching the filter criteria (only.pos = TRUE, min.pct = 0.25, logfc.threshold = 0.25) were kept.

#### Correlation analysis

For the purpose of comparing gene expression levels with the scRNA-seq data from FFPE and fresh sample, we applied standard normalization and scaling procedures. The average normalized expression values were calculated using Seurat’s “AverageExpression” function. The natural logarithm of the average expression with one added pseudo count was plotted and the coefficient of variation and *p*-value was calculated using ggpubr package (0.4.0) in R.

#### Functional Pseudotime Analysis

The differentiation trajectory of a set of glioma cells was performed with Monocle3 and Seurat packages in R.

#### Analyses of lipid metabolic pathways

The visualization and quantifying the lipid metabolic diversity of single cells in each cluster were performed with scMetabolism (v0.2.1)^53^ package in R.

#### Group preference analysis

Group preference of each cell states in primary-recurrent GBM samples was calculated by Chi-Square test (*R*_O/E_)^54^ using chisq.test function in R.

## Data availability

The AAsnRandom-seq data generated in this study have been deposited in the Genome Sequence Archive database. Source data are provided with this paper.

## Code availability

The code for the preprocessing of AAsnRandom-seq data is available at https://github.com/wanglab2023/smRandom-seq.

## Acknowledgements

The project was supported by the National Natural Science Foundation of China (No. 82200977, No. 32200073), Leading Innovative and Entrepreneur Team Introduction Program of Zhejiang (No. 2021R01012), Beijing Xisike Clinical Oncology Research Foundation Y-zai2021/qn-0204 and Y-zai2021/zd-0207. Thanks for the technical support by the core facilities of Zhejiang University and Liangzhu Laboratory. We thank Jingyao Chen and Chengcheng Zhang from the Core Facilities, Zhejiang University School of Medicine for their technical support.

## Author contributions

Y.W., Z.Q., Q.L., and Y.S. conceived the study and managed the project progress. Y.W., and Z.X. coordinated the experiments and analysis. Y.L., M.Z., H.Z., and Q.L. constructed the automated and AI-assistant platform. Z.X. and H.C. developed the AAsnRandom-seq chemistry and performed experiments. Z.X., L.C., X.L., and T.Z. performed data analysis and visualization. L.C., X.L., Y.C., and Z.Q. collected samples and clinical information. Other authors assisted with the experiments and participated in critical discussions. Z.X. and Y.L. wrote the paper. All authors have revised and approved the final manuscript.

## Supplementary information

Supplementary Figure 1-9

Supplementary Table 1

Supplementary Table 2

