## Supplementary information for "An automated archival single-nucleus total RNA sequencing platform mapping integrative and retrospective cell atlas of gliomas"

Supplementary Figure 1

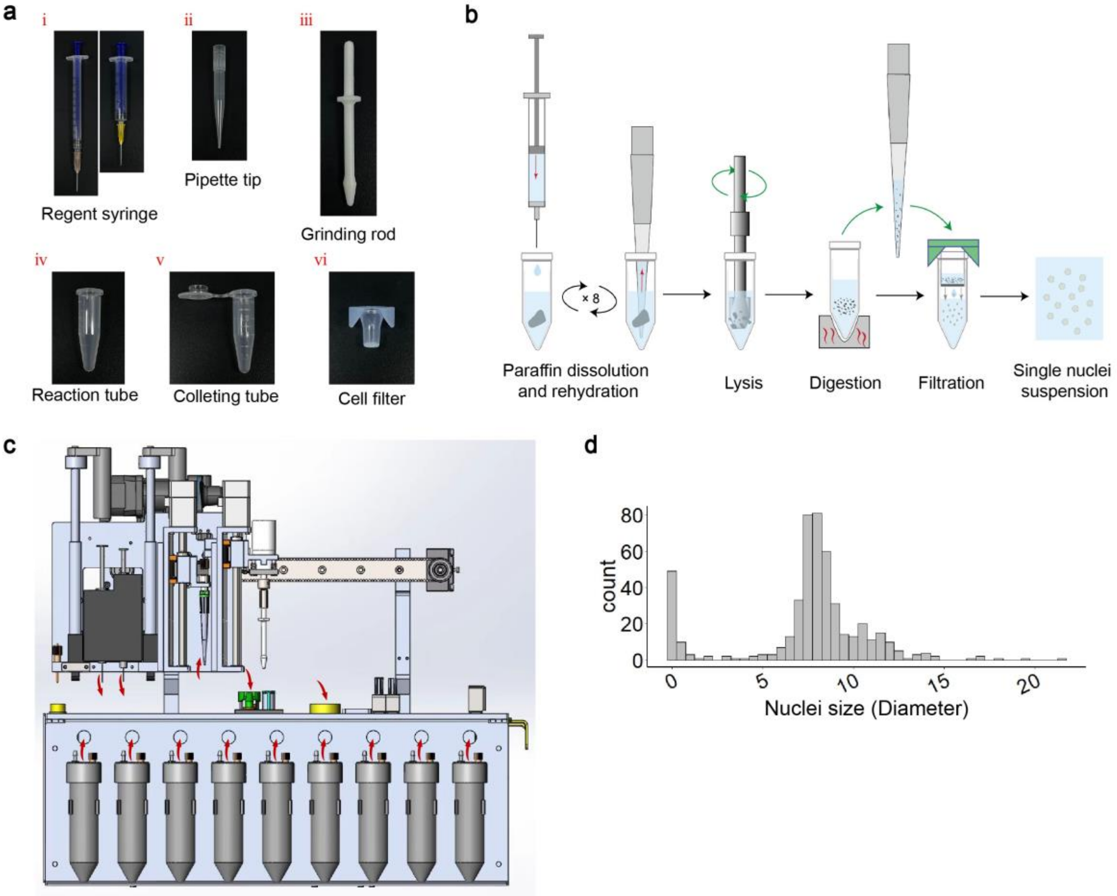


**Supplementary Figure 1. Automated single-nucleus isolation system within AAsnRandom-seq.**

**a**, Photographs depicting the accessories used in the automated single nucleus isolation system of AA-snRandomseq. **b**, A schematic diagram outlining the workflow of the automated single nucleus pretreatment system, including four stages: paraffin dissolution and rehydration (exclusive to FFPE samples), lysis, digestion, and filtration. **c**, A schematic diagram showed the internal structure and flow directions within the automated pretreatment system. **d**. Histograms depicting the distribution of single nucleus diameters. Individual number: 6. Source data are provided as a Source Data file.

Supplementary Figure 2

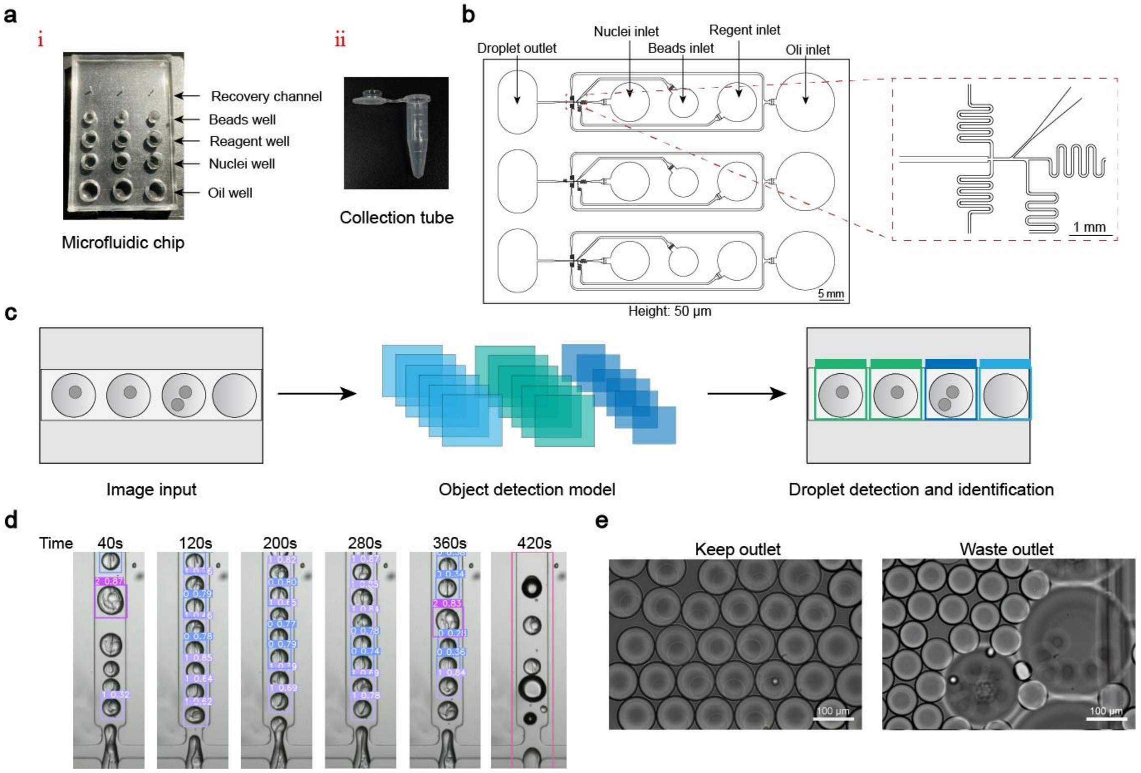


**Supplementary Figure 2. Automated droplet barcoding system within AAsnRandom-seq.**

**a,** Photographs depicting of the accessories used in the droplet barcoding platform of AA-snRandomseq. **b**, Design of the device for the automated encapsulation of cells, beads and mix. Height: 50 µm. Scale bar: 1 mm. Arrows indicate the functions of the holes. A red dashed box indicates a local enlargement. **c**, A schematic illustrating the YOLOv5 object detection model for droplet detection and identification. **d**, Screen images displaying the results of droplet identification at 40, 120, 200, 280, 360, 420 seconds during the automated droplet encapsulation process. **e**, Microscope images showing droplets collected in keeping (left panel) and waste (right panel) tubes. Scale bar: 100 µm.

Supplementary Figure 3

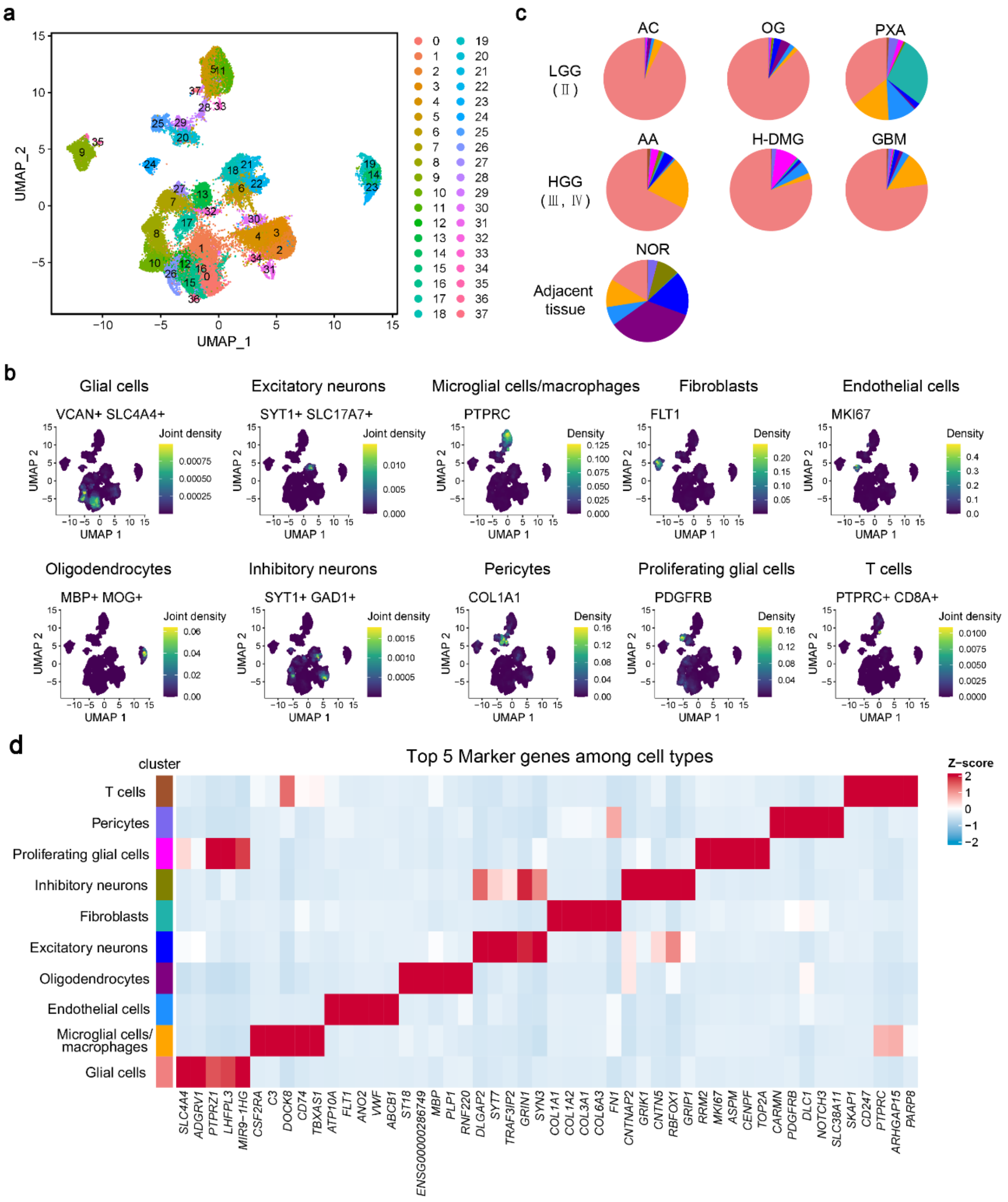


**Supplementary Figure 3. Single-nuclei transcriptome characteristics of gliomas**

**a,** UMAP analysis of nuclei isolated from the seven FFPE samples by AAsnRandom-seq. UMAP plot is colored by Seurat clusters. **b**, Visualization of the expression of marker genes for cell populations on the UMAP dimensions using the kernel density function (plot_density). **c**, Pie charts illustrating the proportion of each cell cluster in each of the applied samples. LGG: low-grade glioma, HGG: high-grade glioma. **d**, Heatmap displaying the expression of top 5 marker genes in each cell type.

Supplementary Figure 4

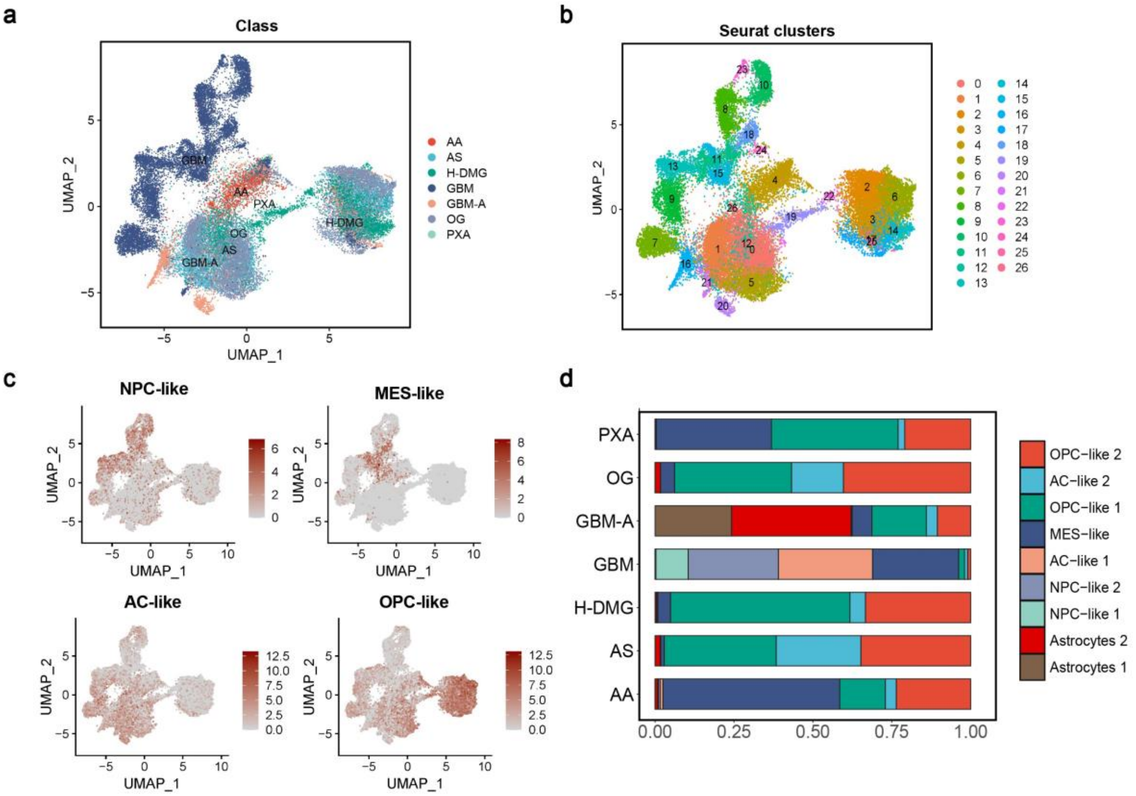


**Supplementary Figure 4. Single-nucleus transcriptome characteristics of glia cells**

**a, b,** UMAP analysis of glia cells from the seven FFPE samples by AAsnRandom-seq, colored by sample classes (**a**) and Seurat clusters (**b**). **c**, Visualization of marker gene expressions for the four states of glial cells on the UMAP dimensions using the kernel density function (plot_density). NPC-like cluster: *STMN2*, *DLX2*, *SATB2*, and *PGM2L1*. MES-like cluster: *CHI3L1* and *ADM*. AC-like cluster: *GFAP*, *SOX9*, *HOPX*, and *HEPACAM*. OPC-like cluster: *OLIG1*, *TNR*, and *ALCAM*. **d,** The percentage bar chart showing the proportion of different cell states in each applied sample.

Supplementary Figure 5

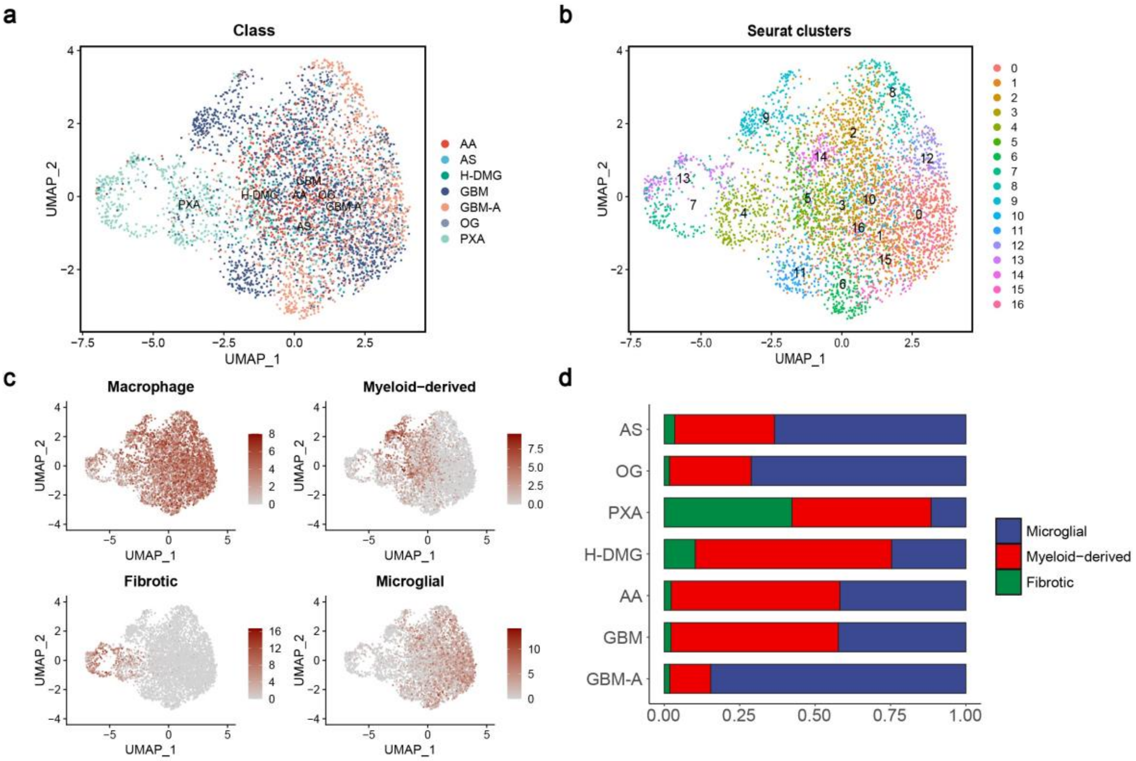


**Supplementary Figure 5. Single-nucleus transcriptome characteristics of microglial cells/macrophages**

**a, b,** UMAP analysis of macrophages from the seven FFPE samples by AAsnRandom-seq, colored by sample classes (**a**) and Seurat clusters (**b**). **c**, Visualization of marker gene expressions for various microglial cell/macrophage subtypes on the UMAP dimensions using the kernel density function (plot_density). Macrophage cluster: *PTPRC* and *CSF1R*. Fibrotic cluster: *COL1A1*, *COL1A2*, *COL6A3*, and *COL6A2*. Myeloid-derived cluster: *CD163*, *TGFBI*, and *F13A1*. Microglial cluster: *CX3CR1*, *P2RY12*, *P2RY13*, and *SELPLG*. **d,** The percentage bar chart showing the proportion of different microglial cell/macrophage subtypes in each applied sample.

Supplementary Figure 6

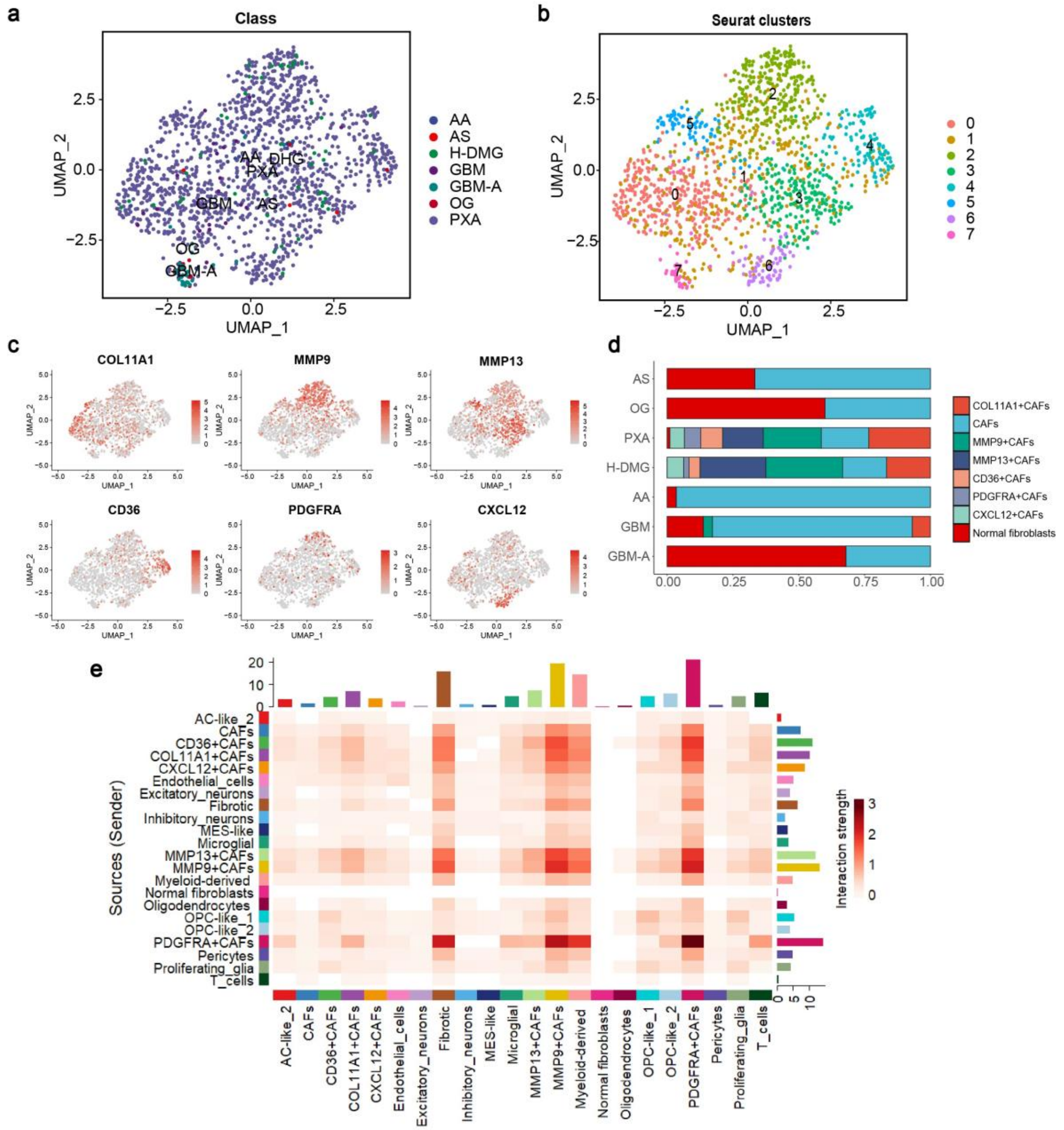


**Supplementary Figure 6. Single-nucleus transcriptome characteristics of fibroblasts**

**a, b,** UMAP analysis of fibroblasts from the seven FFPE samples by AAsnRandom-seq, colored by sample classes (**a**) and Seurat clusters (**b**). **c**, Visualization of marker gene expression for distinct fibroblast subtypes on the UMAP dimensions using the kernel density function (plot_density). **d,** Percentage bar chart illustrating the proportion of different fibroblast subtypes in each analyzed sample. **e,** The heatmap shows the strength of cell communications among subclusters in PXA.

Supplementary Figure 7

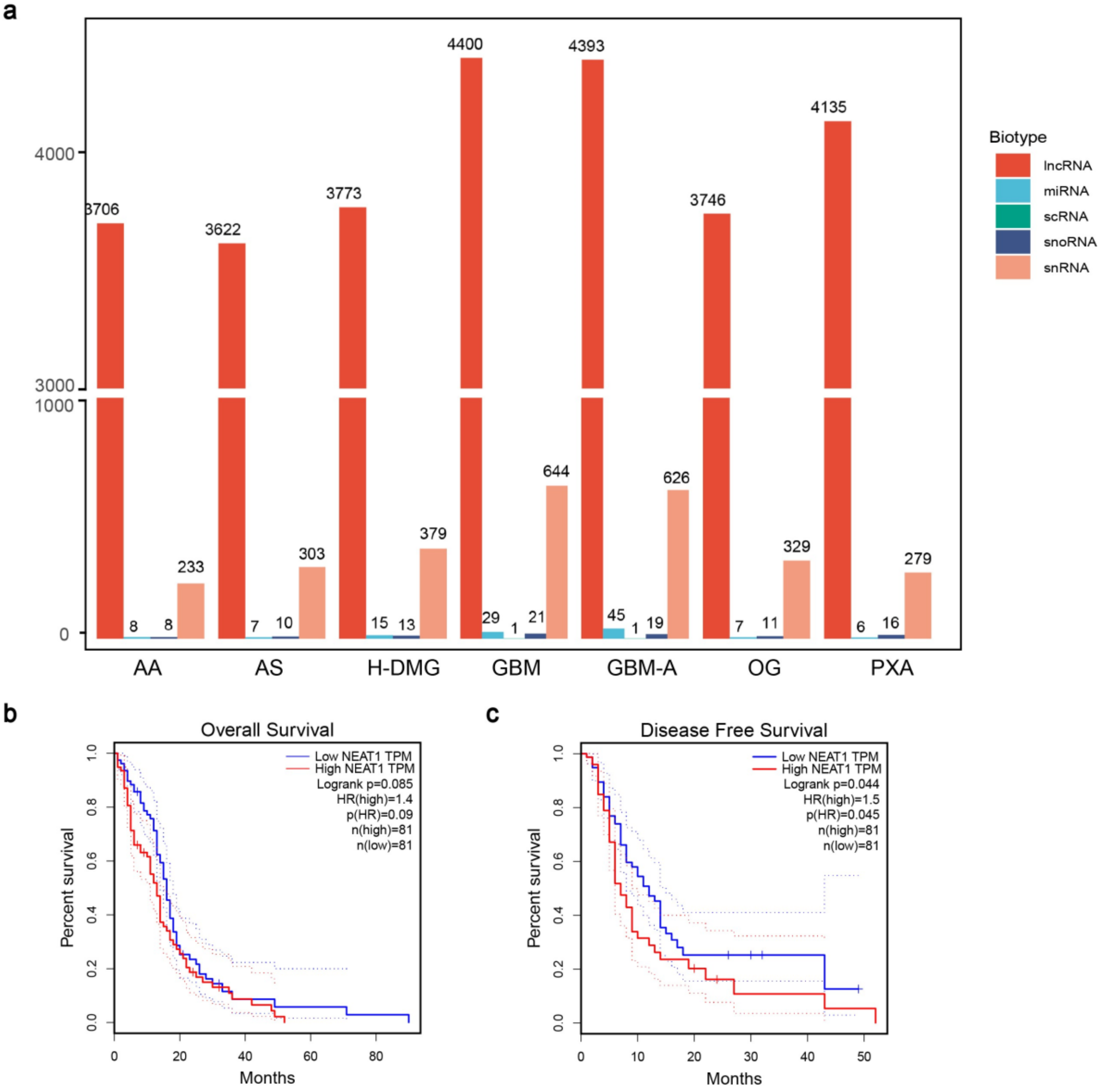


**Supplementary Figure 7. Non-coding RNAs detected by AAsnRandom-seq in glioma FFPE samples. a,** The bar chart showing the counts of lncRNAs, miRNAs, snRNAs, and snoRNAs across all FFPE glioma samples. **b**, **c**, The overall survival (OS) (**b**) and disease free survival (DFS, also called relapse-free survival and RFS) (**c**) analysis of lncRNA *NEAT1* was performed using GEPIA 2 database[1] (http://gepia2.cancer-pku.cn). The glioblastoma (GBM) dataset in GEPIA 2 was selected. Cutoff of high-expression cohort was 50%. Cutoff of low-expression cohort was 50%. The hazards ratio (HR) was calculated based on Cox PH Model. GEPIA uses Log-rank test, a.k.a the Mantel-Cox test, for hypothesis test (*p*(HR)). The dotted lines represent the 95% confidence interval information. The X-Axis units is months.

Supplementary Figure 8

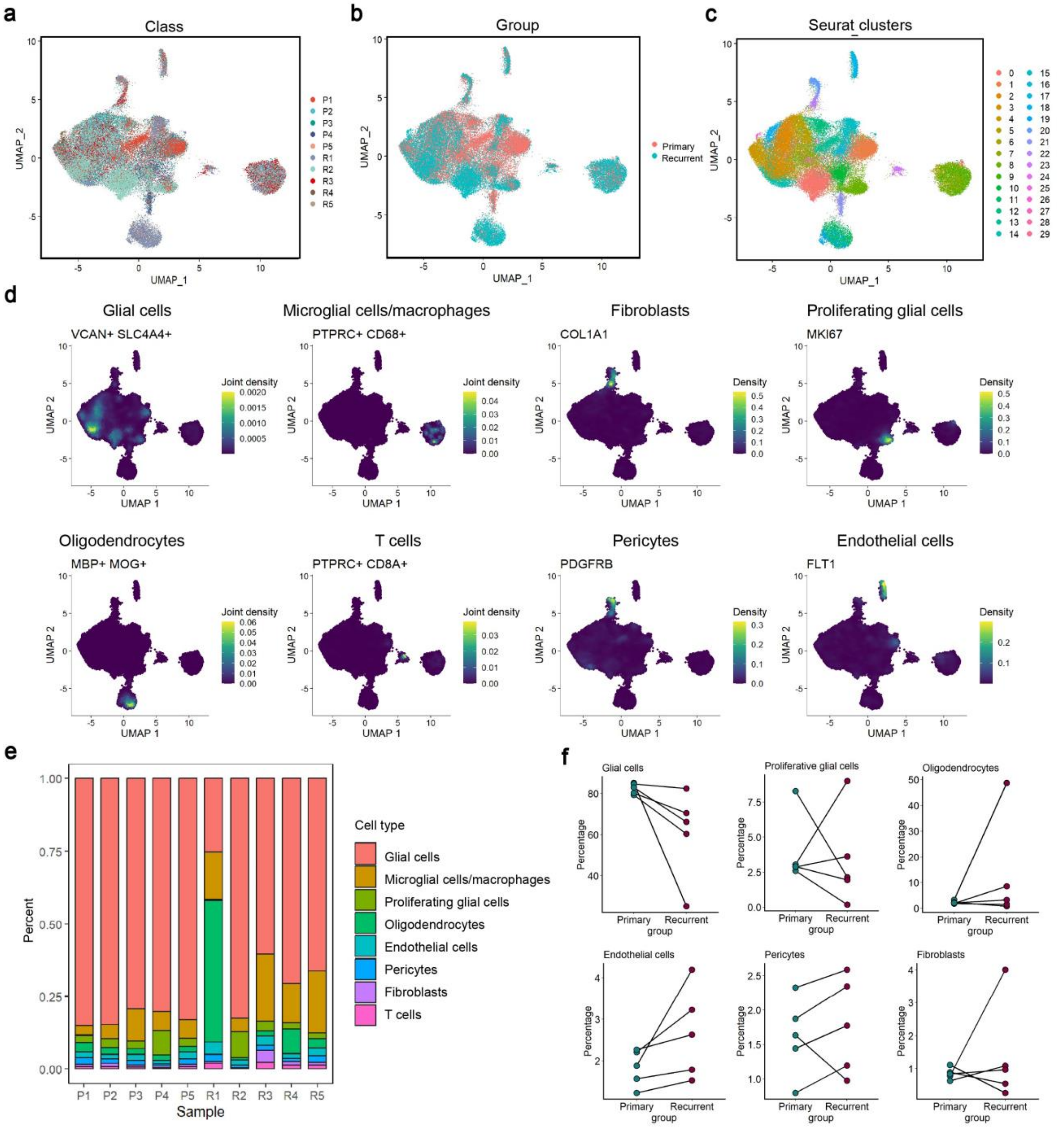


**Supplementary Figure 8. Single-nucleus transcriptome characteristics of matched primary-recurrent GBM samples.**

**a-c,** UMAP analysis of nuclei isolated from five pairs of matched primary-recurrent GBM samples using AAsnRandom-seq, colored by sample classes (**a**), groups (**b**), and Seurat clusters (**c**). **d**, Visualization of marker gene expressions for various cell types on the UMAP dimensions using the kernel density function (plot_density). **e,** The percentage bar chart showing the distribution of different cell types in each applied sample. **f,** The scatter plot showing the percentages of different cell types in each applied sample.

Supplementary Figure 9

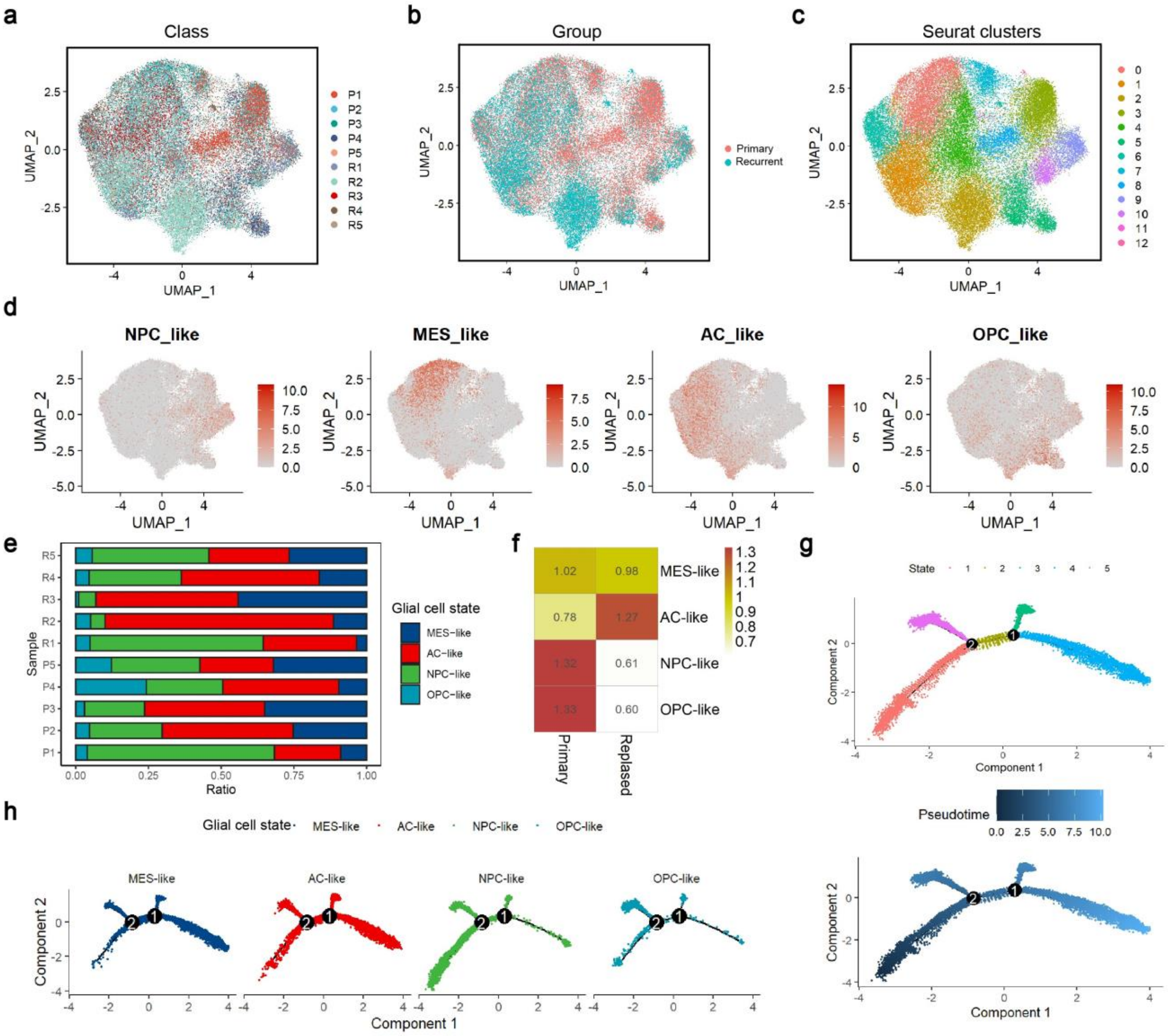


**Supplementary Figure 9. Single-nucleus transcriptome characteristics of glial cells from matched primary-recurrent GBM samples.**

**a-c,** UMAP analysis of glial cells from five pairs of matched primary-recurrent GBM samples by AAsnRandom-seq and colored by samples (**a**), groups (**b**), and Seurat clusters (**c**). **d**, Visualization of marker gene expressions for different glial cell states on the UMAP dimensions using the kernel density function (plot_density). **e,** The percentage bar chart showing the distribution of different glial cell states in each applied sample. **f,** Group preference of each glial cell state in primary and recurrent GBM groups. The ratio of observed to expected cell numbers calculated by the chi-square test ($R_{O/E}$). **g**, Trajectories of glial cells from primary and recurrent GBM samples by monocle analysis and colored by states (upper panel) and pseudotime (lower panel). **h**, Separated trajectories of glial cells from primary and recurrent GBM samples by monocle analysis and colored by glial cell states.

Supplementary Table 1

Supplementary Table 1: Clinical samples applied with AAsnRandom-seq

| Case | System | Lysis buffer | Digestive enzyme | Digestion time (min) | Case number |
|---|---|---|---|---|---|
| Heart muscle | Circulatory system | High | Protein K | 30 | 2 |
| Heart valve | Circulatory system | High | Protein K | 30 | 2 |
| Infantile hemangioma | Circulatory system | High | Protein K | 20 | 3 |
| Gastric carcinoma | Digestive system | Low | Collagenase+ Protein K | 45+ 10 | 2 |
| Cholangiocarcinoma | Digestive system | High | Protein K | 15 | 3 |
| Liver cancer | Digestive system | High | Protein K | 30 | 12 |
| Pancreatic cancer | Digestive system | High | Protein K | 25 | 1 |
| Colorectal cancer | Digestive system | Low | Collagenase+ Protein K | 45+ 10 | 20 |
| Parotid gland carcinoma | Digestive system | High | Protein K | 15 | 2 |
| Liposarcoma | Endocrine system | High | Protein K | 15 | 1 |
| Brain lymphoma | Immune system | Medium | Protein K | 15 | 3 |
| Lymphoma | Immune system | Medium | Protein K | 15 | 2 |
| Rhabdomyosarcoma | Motor system | High | Protein K | 30 | 4 |
| Glioma | Nervous system | High | Protein K | 30 | 25 |
| Brain metastasis of lung cancer | Nervous system | High | Protein K | 25 | 1 |
| Cerebral vasculitis | Nervous system | High | Protein K | 25 | 2 |
| Hypophysoma | Nervous system | High | Protein K | 15 | 2 |
| Ovarian cancer | Reproductive system | Medium | Protein K | 15 | 7 |
| Breast cancer | Reproductive system | High | Protein K | 15 | 6 |
| Cervical cancer | Reproductive system | Medium | Protein K | 20 | 1 |
| Endometrial cancer | Reproductive system | Medium | Protein K | 25 | 1 |
| Lung cancer | Respiratory system | Medium | Collagenase+ Protein K | 45+10 | 21 |
| Nasopharyngeal cancer | Respiratory system | High | Protein K | 30 | 6 |
| Total | | | | | 129 |

Supplementary Table 2

Supplementary Table 2: Clinical and genomic characteristics of glioma samples.

| Patient | Sample | Tumor type | WHO Grade | Gender | Age | Position | Pathology | IDH1 | GFAP | Olig2 | P53 | ATRK | H3K27ME | H3K27M |
|---|---|---|---|---|---|---|---|---|---|---|---|---|---|---|
| G1 | Normal | / | / | Female | 34 | Left frontal corpus callosum | Normal | / | / | / | / | / | / | / |
| | GBM | Glioblastoma | IV | | | | Glioblastoma | - | / | / | / | / | / | / |
| G2 | AG | Astroglioma | II | Female | 45 | Right temporal lobe | Astroglioma | + | + | - | + | + | - | - |
| G3 | ODG | Oligodendroglioma | II | Female | 56 | Right frontal lobe | Oligodendroglioma | + | + | + | - | + | + | - |
| G4 | PXA | Pleomorphic xanthoastrocytoma | II | Female | 26 | Left temporal lobe | Astroglioma | - | + | + | + | + | + | - |
| G5 | H-DMG | High-grade diffuse midline gliomas | IV | Female | 48 | Mesial line of right parietal lobe | Diffuse midline glioma | - | + | + | + | + | - | + |
| G6 | AA | Anaplastic astrocytoma | III | Male | 63 | Right thalamus | Astroglioma | - | + | + | + | + | + | + |
| G7 | R1 | Glioblastoma | IV | Male | 48 | Right parieto-occipital lobe | Glioblastoma | - | + | + | - | + | + | - |
| | P1 | Glioblastoma | | | 46 | Right basal ganglia | Glioblastoma | - | + | -/+ | + | + | + | - |
| G8 | R2 | Glioblastoma | IV | Female | 63 | Left top pillow, left question | Glioblastoma | - | + | + | + | + | + | - |
| | P2 | Glioblastoma | | | 63 | Left frontoparietal | Glioblastoma | - | + | + | + | + | + | - |
| G9 | R3 | Glioblastoma | IV | Female | 51 | Right frontal lobe | Glioblastoma | - | + | + | + | + | + | - |
| | P3 | Glioblastoma | | | 50 | Right frontal lobe | Glioblastoma | - | + | + | + | +/- | -/+ | - |
| G10 | R4 | Glioblastoma | IV | Male | 59 | Right frontal parietal lobe | Glioblastoma | - | + | + | + | + | + | - |
| | P4 | Glioblastoma | | | 59 | Right frontal parietal lobe | Glioblastoma | - | + | + | + | -/+ | -/+ | - |
| G11 | R5 | Glioblastoma | IV | Male | 49 | Right frontal lobe | Glioblastoma | - | + | + | + | + | + | - |
| | P5 | Glioblastoma | | | 49 | Right occipital lobe | Glioblastoma | - | + | + | + | + | + | - |
